# A Knowledge-guided Mechanistic Model of Synthetic Lethality in the HCT116 Vorinostat-resistant Colon Cancer Xenograft Model Cell-line

**DOI:** 10.1101/2021.06.22.449530

**Authors:** Paul Aiyetan

## Abstract

With an overall lifetime risk of about 4.3% and 4.0%, in men and women respectively, colorectal cancer remains the third leading cause of cancer-related deaths in the United States. In persons aged 55 and below, its rate increased at 1% per year in the years 2008 to 2017 despite the steady decline associated with improved screening, early diagnosis and treatment in the general population. Besides standardized therapeutic regimen, many trials continue to evaluate the potential benefits of vorinostat, mostly in combination with other anti-neoplastic agents for its treatment. Vorinostat, an FDA approved anti-cancer drug known as suberoylanilide hydroxamic acid (SAHA), an histone deacylase (HDAC) inhibitor, through many mechanisms, causes cancer cell arrest and death. However, like many other anti-neoplastic agents, resistance and or failures have been observed. In the HCT116 colon cancer cell line xenograft model, exploiting potential lethal molecular interactions by additional gene knockouts restored vorinotat sensitivity. This phenomenon, known as synthetic lethality, offers a promise to selectively target cancer cells. Although without clearly delineated understanding of underlying molecular processes, it has been demonstrated as an effective cancer-killing mechanism. In this study, we aimed to elucidate mechanistic interactions in multiple perturbations of identified synthetically lethal experiments, particularly in the vorinostat-resistant HCT116 (colon cancer xenograft model) cell line. Given that previous studies showed that knocking down GLI1, a downstream transcription factor involved in the Sonic Hedgehog pathway – an embryonal gene regulatory process, resulted in restoration of vorinostat sensitivity in the HCT116 colorectal cancer cell line, we hypothesized that vorinostat resistance is a result of upregulation of embryonal cellular differentiation processes; we hypothesized that elucidated regulatory mechanism would include crosstalks that regulate this biological process. We employed a knowledege-guided fuzzy logic regulatory inference method to elucidate mechanistic relationships. We validated inferred regulatory models in independent datasets. In addition, we evaluated the biomedical significance of key regulatory network genes in an independent clinically annotated dataset. We found no significant evidence that vorinostat resistance is due to an upregulation of embryonal gene regulatory pathways. Our observation rather support a topological rewiring of canonical oncogenic pathways around the PIK3CA, AKT1, RAS/BRAF etc. regulatory pathways. Reasoning that significant regulatory network genes are likely implicated in the clinical course of colorectal cancer, we show that the identified key regulatory network genes’ expression profile are able to predict short- to medium-term survival in colorectal cancer patients – providing a rationale basis for prognostification and potentially effective combination of therapeutics that target these genes along with vorinostat in the treatment of colorectal cancer.

## Introduction

The quest for effective therapies for colorectal cancer, particular in younger patients with advanced disease has never been more imperative. With an overall lifetime risk of approximately 4.3% and 4.0%, in men and women respectively[1, 2], colorectal cancer is the second leading cause of cancer-related deaths in the United States[3]. In persons aged 50 and below, its rate increased at 2% per year in the years 2012 to 2016 despite the steady decline associated with improved screening, early diagnosis and treatment in the general population[2, 3]. According to the center for disease control and prevention (CDC), in 2017, 141, 425 new cases of colorectal cancers were reported, and 52, 547 people died of it[4]. The CDC estimates that for every 100, 000 people, 37 new colorectal cancer cases are reported and 14 people died of this cancer[4].

Historically, risk factors have been classified as modifiable and non-modifiable factors[5]. Modifiable factors have included being overweight, a sedentary lifestyle, diet rich in red and processed meat, and sugars, smoking and alcohol consumption, while non-modifiable factors include increasing age, history of inflammatory bowel disease, polyps, family history of colorectal cancer, ethnicity, type II diabetes mellitus, and familial or inherited syndromes [5]. Although familial or hereditary factors account for only a third of colorectal cancer diagnoses, their molecular basis have enabled fundamental understanding of the etiopathogenesis of the disease. These include, lynch syndrome (hereditary non-polyposis colon cancer or HNPCC) which is primarily associated with defects in the *MLH1, MSH2* or the *MSH6* genes, and accounts for about 2% to 4% of all colorectal cancers, familial adenomatous polyposis coli (FAP) which accounts for 1% of colorectal cancers, Peutz-Jeghers syndrome (PJS), and MUTYH-associated polyposis (MAP). Associated with mutations in the APC gene, the FAP-related colorectal cancer consists of three sub-types with almost specific clinical features. These include: the attenuated FAP, associated with fewer polyps and development of colorectal cancer at a later age than it is typical; the Gardner syndrome, associated with tumors of the soft tissues, bones and skin; and the Turcot syndrome, associated with an higher risk of colorectal cancer and a predisposition to developing medulloblastoma – a brain cancer. Usually diagnosed at a younger age, PJS is associated with mutations in the *STK11* (*LKB1*) gene while as its name implies, MAP is caused by mutations in the *MUTYH* gene[5]. These associated genetic defects are characteristically those of genes involved in tumor suppression and DNA repair mechanisms [6].

Besides standardized therapeutic regimen, many trials continue to evaluate the potential benefits of vorinostat, mostly in combination with other anti-neoplastic agents for its treatment[7–17]. Vorinostat, an FDA approved anti-cancer drug known as suberoylanilide hydroxamic acid (SAHA), a histone deacetylase (HDAC) inhibitor, through many mechanisms, causes cancer cell arrest and death[18]. First discovered on attempts to make more efficient hybrid polar compounds that induce the differentiation of transformed cells[19, 20] and initially approved by the FDA for the cutanous manifestation of T cell leukemia, vorinostat has since become a therapeutic candidate for many tumors[21–29]. This is due in part to the evolving understanding of the role of epigenetic and posttranslational modifications in the etiopathogenesis of transformed cells[30–33]. Altering many pathways and processes, vorinostat has been discovered to not only alter the modification state of histone proteins but many more essential proteins involved in the oncogenic and tumor suppression process. More specifically and among many other mode of action, vorinostat inhibits the removal of acetyl group from the *ϵ*-amino group of lysine residues of histone proteins by histone deacetylases (HDACs). Accumulation of acetyl group maintains chromatin in an expanded state, facilitating transcriptional activities of major regulatory genes[18, 30, 34–36]. However, like many other anti-neoplastic agents, toxicities, resistance and or failures have been observed[13, 37, 38].

In the HCT116 colon cancer cell line xenograft model, exploiting potential lethal molecular interactions by additional gene knockouts, Falkenberg and colleagues were able to restore vorinotat sensitivity[39, 40]. This phenomenon, known as synthetic lethality, offers a promise to selectively target cancer cells[41]. Although without clear delineated understanding of underlying molecular processes, many studies demonstrate synthetic lethality as an effective cancer-killing mechanism.

In this study, we aimed to elucidate regulatory interactions, in multiple perturbations of identified synthetically lethal experiments, particularly in the vorinostat-resistant HCT116 (colon cancer xenograft model) cell line. In addition to elucidating interactions, we aim to elucidate key interactions that potentially determine observed phenotypes. Given that previous studies[39, 40] showed that knocking down GLI1, a downstream transcription factor involved in the Sonic hedgehog (SHH) pathway [42–44] – an embryonal gene regulatory process, resulted in restoration of vorinostat sensitivity in the HCT116 colorectal cancer cell line, we hypothesized that vorinostat resistance is a result of uptick in embryonal gene regulatory programs. We also hypothesized that elucidated regulatory mechanism would include crosstalks that regulate this biological processes – embryonal gene regulatory programs. We employed a knowledege-guided fuzzy logic regulatory inference method to elucidate mechanistic relationships from multiple synthetic lethal pertubation experiments in the vorinostat-resistant colon cancer cell lines. We validated inferred regulatory models in independent experiment datasets. And, we evaluated the biomedical significance of key regulatory network genes in an independent clinically annotated dataset.

## Materials and Methods

### Datasets

#### Synthetic Lethal Experiments Transcriptome, RNA Sequencing Assay Data

Two RNASeq expression datasets (accessions GSE56788 and GSE57871) with available viability assay data were retrieved from the National Center for Biotechnology information (NCBI) Gene Expression Omnibus [45, 46] public repository.

#### GSE56788

Detailed under the BioProject accession PRJNA244587, this consists of 45 assays from 15 biosamples, each ran in 3 independent biological replicates. RNA-seq expression profiles were acquired by next-generation sequencing of vorinostat-resistant HCT116 cells (HCT116-VR) following knockdown of potential vorinostat-resistance candidate genes. Assays included those of mock transfection to serve as controls. The authors of the study sought to understand the mechanisms by which these knockdowns contributed to vorinostat response – reestablishment of a gain in sensitivity to Vorinostat. siRNA-mediated knockdown of each previously identified resistance candidate genes in the HCT116-VR cell line was employed[39]. Raw RNA sequence expression data were downloaded from the NCBI Sequence Read Archive [47, 48], with accession number SRP041162. Table 1 shows the transcriptome expression profile data accessions and associated siRNA treatment experiments.

**Table 1:**
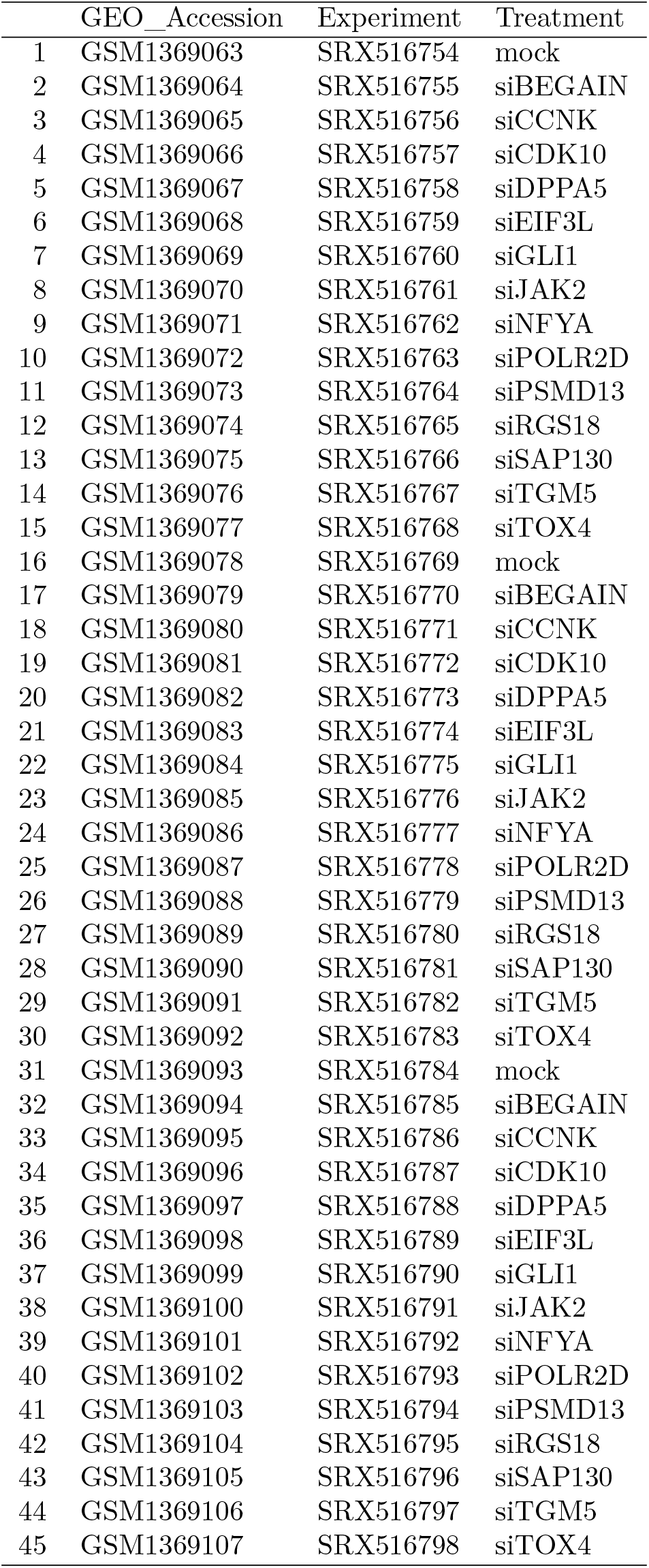
GSE56788 Gene Expression Omnibus, GEO dataset I

#### GSE57871

Similar to GSE56788, the GSE57871 study is a 42 sample dataset derived from an expression profiling by high throughput sequencing. It consists of independent biological experiments of 14 samples performed in triplicates. RNA-seq high throughput expression profiling of vorinostat-resistant HCT116 cells was performed following gene knockdown of GLI1 or PSMD13 with or without vorinostat treatment. Study authors had chosen GLI1 and PSMD13 as potential vorinostat resistance genes because these had previously been identified through a genome-wide synthetic lethal RNA interference screen (the GSE56788 dataset study). An aim was to understand the transcriptional events underpinning the effect of GLI1 and PSMD13 knockdown (sensitisation to vorinostat-induced apoptosis). The authors first performed a knockdown on cells, and then treated these with vorinostat or the solvent control. Two timepoints for drug treatment were assessed: a time-point before induction of apoptosis (4hrs for siGLI1 and 8hrs for siPSMD13) and a timepoint when apoptosis could be detected (8hrs for siGLI1 and 12hrs for siPSMD13)[40]. Raw sequence expression data were downloaded from the NCBI Sequence Read Archive with accession number SRP042158. Table 2 shows the transcriptome expression profile sample data accessions and associated siRNA treatment and treatment timepoint experiments.

**Table 2:** GSE57871 Gene Expression Omnibus, GEO dataset

|  | GEO_Accession | Experiment | Timepoint | siRNARx | DrugRx |
| --- | --- | --- | --- | --- | --- |
| 1 | GSM1395357 | SRX548949 | 4hr | mock | DMSO |
| 2 | GSM1395358 | SRX548950 | 8hr | mock | DMSO |
| 3 | GSM1395359 | SRX548951 | 12hr | mock | DMSO |
| 4 | GSM1395360 | SRX548952 | 4hr | mock | vorinostat |
| 5 | GSM1395361 | SRX548953 | 8hr | mock | vorinostat |
| 6 | GSM1395362 | SRX548954 | 12hr | mock | vorinostat |
| 7 | GSM1395363 | SRX548955 | 4hr | siGLI1 | DMSO |
| 8 | GSM1395364 | SRX548956 | 8hr | siGLI1 | DMSO |
| 9 | GSM1395365 | SRX548957 | 4hr | siGLI1 | vorinostat |
| 10 | GSM1395366 | SRX548958 | 8hr | siGLI1 | vorinostat |
| 11 | GSM1395367 | SRX548959 | 8hr | siPSMD13 | DMSO |
| 12 | GSM1395368 | SRX548960 | 12hr | siPSDM13 | DMSO |
| 13 | GSM1395369 | SRX548961 | 8hr | siPSMD13 | vorinostat |
| 14 | GSM1395370 | SRX548962 | 12hr | siPSMD13 | vorinostat |
| 15 | GSM1395371 | SRX548963 | 4hr | mock | DMSO |
| 16 | GSM1395372 | SRX548964 | 8hr | mock | DMSO |
| 17 | GSM1395373 | SRX548965 | 12hr | mock | DMSO |
| 18 | GSM1395374 | SRX548966 | 4hr | mock | vorinostat |
| 19 | GSM1395375 | SRX548967 | 8hr | mock | vorinostat |
| 20 | GSM1395376 | SRX548968 | 12hr | mock | vorinostat |
| 21 | GSM1395377 | SRX548969 | 4hr | siGLI1 | DMSO |
| 22 | GSM1395378 | SRX548970 | 8hr | siGLI1 | DMSO |
| 23 | GSM1395379 | SRX548971 | 4hr | siGLI1 | vorinostat |
| 24 | GSM1395380 | SRX548972 | 8hr | siGLI1 | vorinostat |
| 25 | GSM1395381 | SRX548973 | 8hr | siPSMD13 | DMSO |
| 26 | GSM1395382 | SRX548974 | 12hr | siPSDM13 | DMSO |
| 27 | GSM1395383 | SRX548975 | 8hr | siPSMD13 | vorinostat |
| 28 | GSM1395384 | SRX548976 | 12hr | siPSMD13 | vorinostat |
| 29 | GSM1395385 | SRX548977 | 4hr | mock | DMSO |
| 30 | GSM1395386 | SRX548978 | 8hr | mock | DMSO |
| 31 | GSM1395387 | SRX548979 | 12hr | mock | DMSO |
| 32 | GSM1395388 | SRX548980 | 4hr | mock | vorinostat |
| 33 | GSM1395389 | SRX548981 | 8hr | mock | vorinostat |
| 34 | GSM1395390 | SRX548982 | 12hr | mock | vorinostat |
| 35 | GSM1395391 | SRX548983 | 4hr | siGLI1 | DMSO |
| 36 | GSM1395392 | SRX548984 | 8hr | siGLI1 | DMSO |
| 37 | GSM1395393 | SRX548985 | 4hr | siGLI1 | vorinostat |
| 38 | GSM1395394 | SRX548986 | 8hr | siGLI1 | vorinostat |
| 39 | GSM1395395 | SRX548987 | 8hr | siPSMD13 | DMSO |
| 40 | GSM1395396 | SRX548988 | 12hr | siPSDM13 | DMSO |
| 41 | GSM1395397 | SRX548989 | 8hr | siPSMD13 | vorinostat |
| 42 | GSM1395398 | SRX548990 | 12hr | siPSMD13 | vorinostat |

#### Colon Cancer-Associated Genes from OMIM

A curated list of colon cancer-associated genes (Table 3) were retrieved from the Online Mendelian Inheritance Man (OMIM) database [49, 50].

**Table 3:**
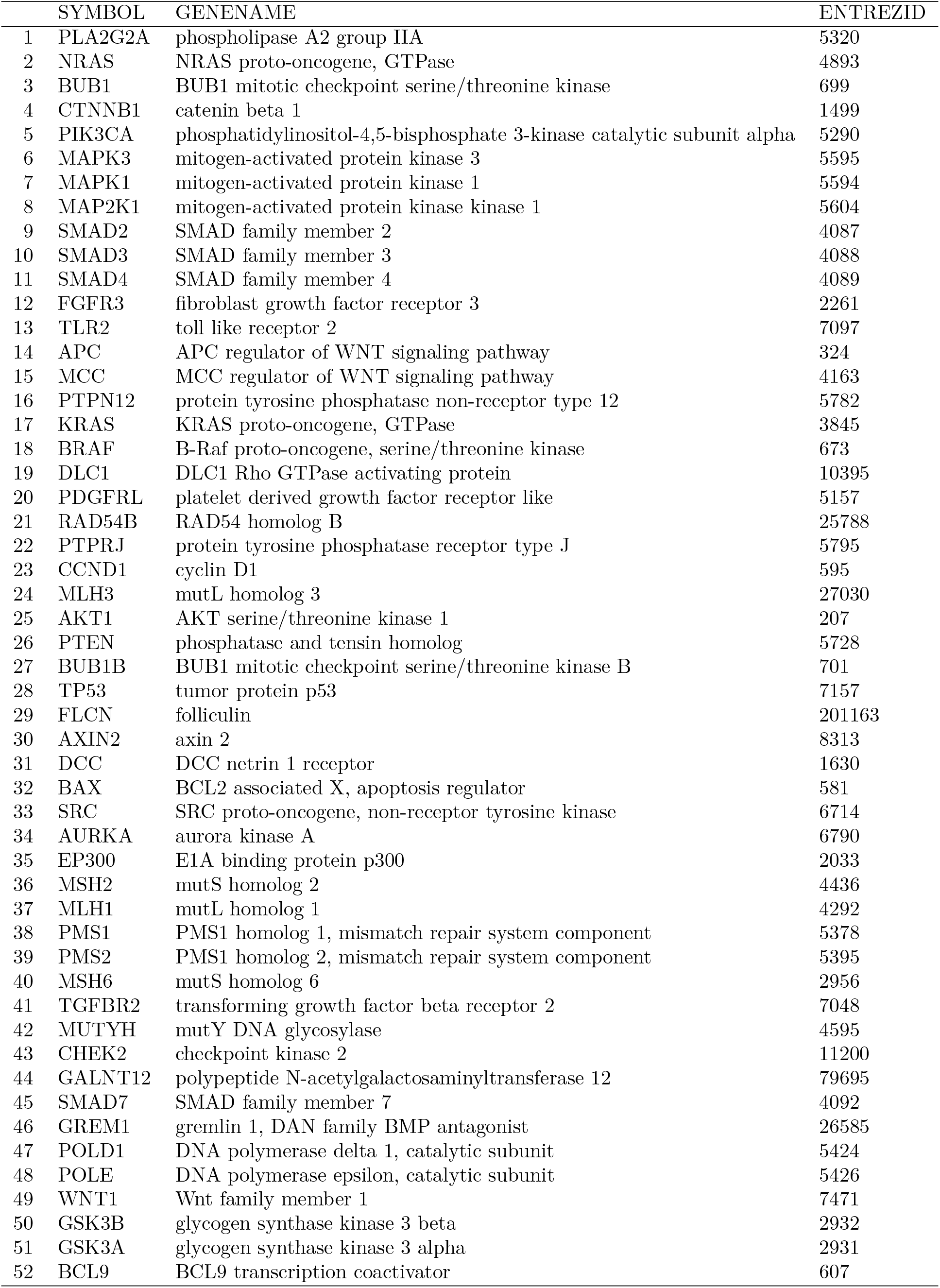
Colon-cancer associated genes

#### Biomedical Significance Experiment Data

To evaluate the clinical and biomedical significance of inferred regulatory features and themes, gene expression profile were retrieved from the cancer genome atlas (TCGA) colorectal cancer mRNA data, in the `TCGAcrcmRNA` R Bioconductor package[51, 52]. The package contains the TCGA consortium-provided level 3 data, generated by the `HiSeq` and `GenomeAnalyzer` platforms, from 450 primary colorectal cancer patient samples[53]. For a more comprehensive and up-to-date phenotype information, associated patients’ clinical data were retrieved from the genomic data commons[54–57].

## Methods

### RNA Sequence Analyses

#### Quality assessment

For data quality assessment (QA), the `fastqcr, ngsReports` and `Rqc` R/bioconductor tools [52, 58–60], modeled after the FASTQC [61] tool philosophy were used. These provide add-on capabilities and the R programming interface to the standalone Java program implementation of FASTQC. QA results were used to identify data with questionable measured quality metrics. In addition to data file statistics, reported quality metrics included; ‘adapter content’, ‘overrepresented sequences’, ‘per base N content’, ‘per base sequence content’, ‘per base sequence quality’, ‘per sequence GC content’, ‘per sequence quality score’, ‘sequence duplication levels’, and ‘sequence length distribution’.

#### Reads quantification

To quantify expression, we aligned reported reads from the sequencing experiment to the genome. Although non-alignment based quantification approaches such as those implemented in Salmon [62], Sailfish [63], and Kallisto [64] are becoming more popular, the performance of these on quantifying lowly expressed genes and small RNAs is still being debated [65]. Therefore sequence reads were aligned to the genome (NCBI GRCh38 build) using the TopHat2 [66, 67] tool which accounts for slice junctions in alignments. Tophat2 uses the bowtie2 [69], noted for its speed and proven memory efficiency for primary alignment. Rather than build new index files, pre-built bowtie2 index files were downloaded from Illumina’s iGenomes archive [70]. Accepted hits and annotation information in the BAM format [71] output files were assembled into an expression matrix of feature counts using the featureCount routine in the Rsubread package [72].

**Figure 1:**
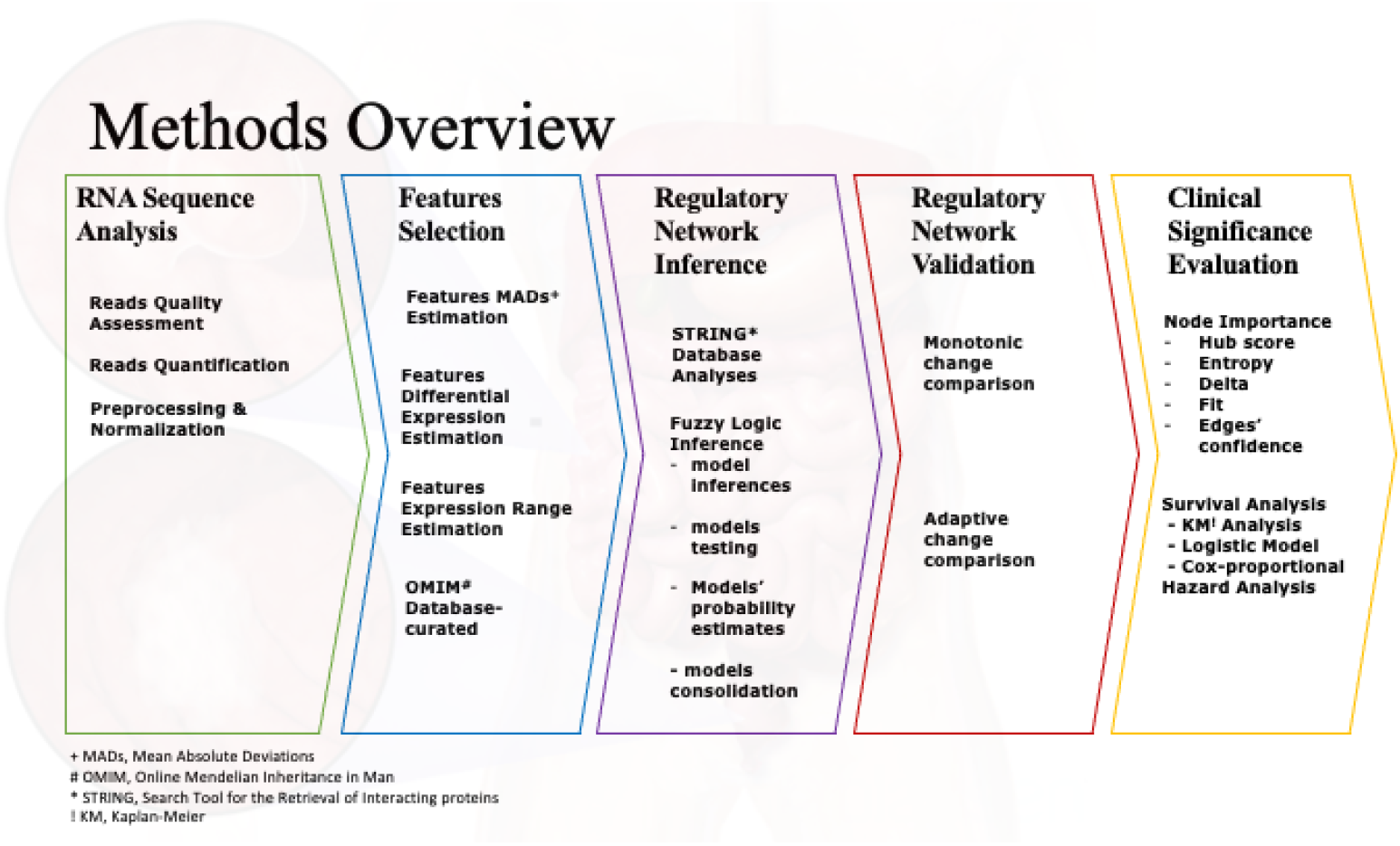
Methods Overview

#### Preprocessing and normalization

Feature counts were normalized using the `DESeq2` package [73] tool’s implemented regularized log transformation to account for disparate total read counts in the different files and to allow for comparison across the different samples. The regularized log transformation moderates the high variance typically observed at low read counts. We specified regularized log transformation intercept as the average expression profile across the normal (mock) samples.

### Model Building and Independent Validation Datasets

Datasets were divided into training (regulatory-model-infering) and test (regulatory-model-validation) datasets (Figure 2). Regulatory models were inferred using the training datasets. Inferred models were tested in the independent validation datasets. Independent validation dataset included two parts. A part was used to test the regulatory models while the other part was used to test and evaluate a simulation of the consolidated network.

**Figure 2:**
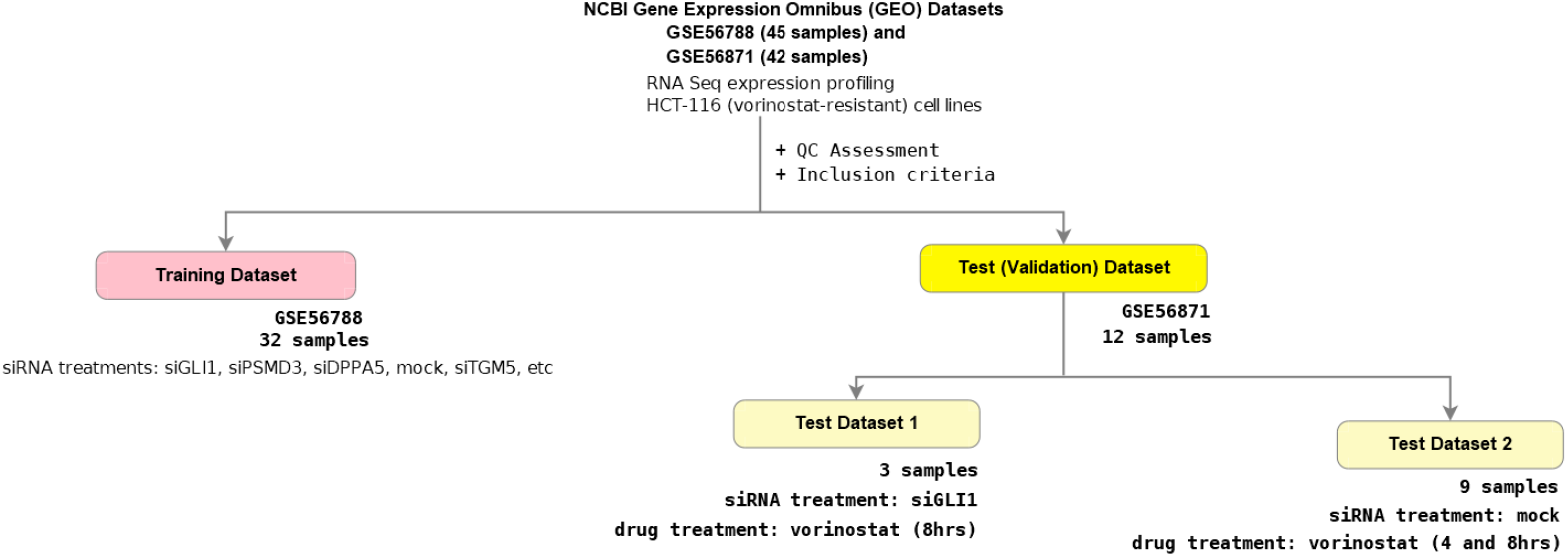
Datasets. For our fuzzy-logic inference and evaluation, Two qualifying datasets, with accession numbers GSE56788 and GSE56871, were found and retrieved from the NCBI Gene Expression Omnibus (GEO) database. The studies’ samples were subjected to quality assessment and inclusion criteria. 32 qualifying samples from the GSE56788 dataset were used for training (model building) and 12 samples meeting our inclusion criteria from the GSE56871 dataset were used for testing. Of the 12 samples, 3 samples from the 12 were derived from GLI siRNA knockdown experiments and 9 samples were from mock experiments.

### Feature Selection

Although similar, feature selection for regulatory network reconstruction and inference differs from classical feature selection. Classical feature selection [74–78] approaches aim to identify the optimal set of features with which a trained model can best predict or correctly identify a class of a not-previously-seen object, given the object’s attributes – the class prediction problem. With a class prediction problem is an associated feature redandancy [79] which needs to be mitigated when choosing an optimal set. With respect to selecting features for regulatory networks however, this may not necessarily be the case, since features that appear redundant may imply co-regulatory (direct or indirect regulatory) interactions in the network. In both situations anyways, on a one hand is the cost of learning a model while on another hand is the curse of dimensionality that plague the low sample to feature ratio characteristic of biological experiments. The very high dimension coupled with low sample size and the potential noise in measured experiments present a limitation for regulatory network inference methods [80] in particular. Feature selection seeks to find a middle spot where cost is minimized with minimal loss in learned model benefits. Although optimized algorithms may mitigate cost, poorly selected or less optimal set of features are set to undermine the efficiency of any learned model.

For a regulatory network model that would represent colon cancer, we reasoned that network features should very likely include known and previously identified products of genes associated with the disease process. Thus, we compiled a list of genes consisting of a curated set obtained from the OMIM database [49, 50] and those from literature evidences i.e. genes in described pathways of colon cancer tumorigenesis. And, if we assume that the regulary network is a function of changes in features’ expression across time, among different perturbations or across cellular states, it should also appeal to reason that features with significant variations or dispersion in expression across samples should be more informative i.e. more relevant for deriving a regulatory network than those without or with minimal variations. Mathematically, we may describe a cellular state *s*, as a linear combination of weighted features’ expressions, given by the equation below:

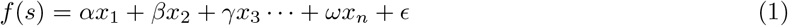

where *α, β, γ*, ··· *ω* are the **rates of change** in respective feature’s expression i.e. *rate constants*; *ϵ* is the random error estimate; {*x*_1_, *x*_2_, *x*_3_ ··· *x_n_*} is the set of expression values of features under condideration; and *n* is the total number of features. We reasoned that if we assume a regulatory network describes changes in cellular state across time, we might as well describe it as a first derivative of cellular state, *f*(*s*)′. Therefore features without changes in expression across time, i.e. features whose rate constants tended to zero would drop off in the estimate *d*(*f*(*s*))/*dt*. This is analogous to being of less significance in determining the dynamic nature of the regulatory network, i.e. changes in cellular state.

To determine maximally varying features, from our RNA sequence analyses normalized expression values, we estimated a mean absolute deviation (MAD) from the mean, for each feature. Given by,

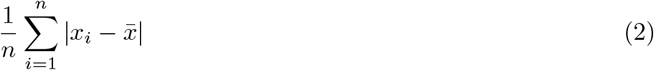

where *n* in this case is the number of samples or perturbations and 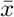 is the mean expression value of the specific feature across the samples. *x_i_* ∈ {*x*_1_, *x*_2_,… *x_n_*}.

To further assess variation in the expression of genes across samples, we also determined fold changes between the minimum and maximum expression values for for the respective genes and the strength of change between knockdown and control experiments. Because genes with highest MADs were observed to be predominantly those with low average expression and thus may be confounded by a Poisson noise distribution, we performed differential expression analyses between the respective groups of knockdown (siRNA) experiments and the controls to identify statistical significantly expressed genes (i.e. features with true changes)[81–83].

In summary, in additon to genes previously identified as related to colon cancer tumorigenesis and the specific genes targeted in the knockdown experiments, expression profile-informed genes were also considered for regulatory network inference based on their MAD, differential expresson and the log fold difference between the minimum and maximum expression values across siRNA knockdown experiments. The expresion profile-based selection criteria we specified were that for a gene to be considered:

1. Its mean absolute deviations (MAD) must be greater than the median of MADs.
2. Its expression value in 80% of samples must be greater than its minimum value across all samples by a minimum of two folds. The 80% of samples must include ≥ 80% of siRNA-targeted experiments. And, it must be
3. Statistical siginificant and differentially expressed in at least two siRNA-targeted sample groups versus the control group

#### Knowledge-guided feature selection

Purely data-driven methods have drawbacks such as limited biological interpretability. Likewise, canonical signaling pathways from literature evidences, provided in curated knowledge databases are not very specific and these hardly predict cell type-specific responses to experimental situations [84]. Therefore, we employed a hybrid approach that addresses these limitations and, can integrate prior knowledge and real data for network inference. We searched the derived features, and the colon cancer related gene features from OMIM database, against the STRING database[85–87]. Our search parameters included: a search against a full network type where edges indicate both functional and physical protein interactions; reported network edges indicate the presence of evidence of interactions between nodes; active interaction sources included mining of literature texts (TextMining), known experiments, knowledge bases, documented co-expression information, gene neighborhood, fusion and co-occurrence information. Quantitative interaction score for retrieved edges was specified as a minimum of 0.150. We retrieved features reported to be part of a potential network. For each feature found as part of a potential network, all reported interacting features were retrieved and mapped. We elaborated regulatory relationships between and among features using the fuzzy logic approach.

### Fuzzy Logic Regulatory Model and Network Inference

To tease regulatory interactions among our initial selection of features, we employed the fuzzy-logic approach. The fuzzy logic approach mitigates known challenges of modeling biological systems, such as inconsistencies and inaccuracies associated with high-throughput characterizations. These challenges also include data noise and those of dealing with a semi-quantitative data [88]. Similar to Boolean networks, fuzzy logic methods are simple and are fit to model imprecise and or highly complex networks. And, opposed to differential equation based models, they are less computationally expensive and less sensitive to imprecise measurements [89–91]. Fuzzy logic compensates for the inadequate dynamic resolution of a Boolean (or discrete) network, while simultaneously addressing the computational complexity of a continuous network [92, 93].

A significant advantage of the fuzzy logic approach is that, in contrast to many other automated decision making algorithms or regulatory inference methods, such as neural networks or polynomial fits, algorithms in fuzzy logic are presented in similar day-to-day conversational language. Therefore, a fuzzy logic is more easily understood and can be extrapolated in predictable ways.

In general, the fuzzy logic modeling approach entails three major steps.

1. Fuzzification
2. Rule evaluation, and
3. Defuzzification

[94].

#### Fuzzification

Considering expression as a linguistic variable and applying defined membership functions on observed continuous numerical expression data, the fuzzification step derives qualitative values. It is a mapping of non-fuzzy inputs to fuzzy linguistic terms [94]. To make data fuzzification easier, a normalization technique may be applied to scale values to within a preferred range [92, 94, 95].

The fuzzification step derives qualitative values from the expression profile’s crisp values. By applying defined membership functions on crisp, numerical expression data, we derived qualitative values – described as a mapping of non-fuzzy inputs to fuzzy linguistic terms [96]. Given qualitative values of HIGH, MEDIUM, or LOW, the fuzzification step takes a feature’s expression value and assigns it degrees to which it belongs to the respective class of HIGH, MEDIUM or LOW expression values. [97–100]. After an initial data transformation of log2 expression ratios by the arctan function and dividing values by 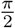, to project the ratios onto [-1,1], the fuzzification step utilizes three membership functions consisting of the ‘low’, ‘medium’, and ‘high’ functions. Given the three fuzzification functions (*y*_1_ = low, *y*_2_ = medium, *y*_3_ = high), fuzzification of a gene expression value *x* results in the generation of a fuzzy set *y* = [*y*_1_, *y*_2_, *y*_3_] as follows:

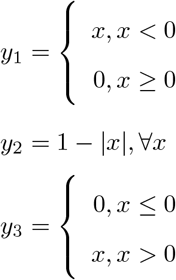

#### Rule evaluation

The rule evaluation step considers combinations of features and utilizes an inference engine of rules, of the form IF-THEN, including fuzzy set operations such as AND, OR, or NOT, to evaluate input features’ expression (in fuzzy set definition) in relation to output features. This has been described as attempting to make an expert judgment of collective linguistic terms; attempts to find a solution to an evaluation of the concurrent state of existence of linguistic description of states.

We specified our rule configuration (the specification of if-then relationships between variables in fuzzy space) in the form of a vector *r* = [*r*_1_, *r*_2_, *r*_3_]. We specified the state of an output node *z* = [*z*_1_, *z*_2_, *z*_3_] to be determined by the fuzzy state of an input feature *y* = [*y*_1_, *y*_2_, *y*_3_] and the rule describing the relationship between the input and the output, *r* = [*r*_1_, *r*_2_, *r*_3_] as follows:

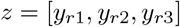

An inhibitory relationship, for example, specified as [3, 2, 1] implies, if input is low (*r*_1_), then output is high (3); if input is medium (*r*_2_), then output is medium (2), and if input is high (*r*_3_), then output in low (1). The classic fuzzy logic rule evaluation using the logical AND connective results in a combinatorial rule explosion i.e. an exponential increase in the number of rules to be evaluated and computational time, with additional inputs to be considered [101]. Therefore, to address this combinatorial rule explosion situation, we employed the logical OR (union) rule configuration, an algebraic sum in fuzzy logic [102, 103] as described in [104].

#### Defuzzification

The defuzzification step produces a quantifiable expression result or value given the input sets, the fuzzy rules, and membership functions. Defuzzification technically interpretes the membership degrees of the fuzzy sets into a specific decision or real value. The defuzzification step attempts to report a corresponding continuous numerical variable from a fuzzy state liguistic variable. Several approaches to defuzzify abound. We employed the simplified centroid method [103]. Given a predicted fuzzy values of an output node *y* = [*y*_1_, *y*_2_, *y*_3_], we defined defuzzified expression values 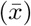 as:

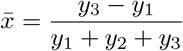

After defuzzification, we reverse transformed back to log2 expression values by multiplying derived values by 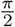 and applying the tangent function.

#### Inferred regulatory model fit

For each regulatory model, which consists of an output feature, its suggested regulatory input feature(s) and associated fuzzy logic rules (relating each input feature to the output respectively), we estimated the fitness of such model’s prediction of the output *x* across *M* experiment samples or perturbations *x* = {*x*_1_, *x*_2_,…, *x_M_*} as:

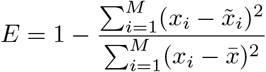

where 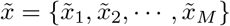 is the set of defuzzified numerical log expression ratios predicted for the output feature and 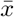 is the mean of the experimental values of *x* across the samples or perturbations observed. A perfect fit would result in a maximum *E* of 1.0.

#### Model probability (p-value) estimates

To estimate models probabilities, we fitted a probability density distribution for 100, 000 fit estimates of models derived by random permutations of rules and input features for each output features. We allowed up to four regulatory interactors. We computed a model fit’s p-value as the probability of observing an estimated fit from a random estimated fits distribution. A gamma distribution was fitted and, the ‘scale’ and ‘shape’ parameters were derived using The Maximum Likelihood Estimate (MLE) approach [105–108] implemented in the egamma function, in the `EnvStat` R package. With the ‘’scale’ and ‘shape’ parameters, random deviates and cummulative probabilities were derived using the (rgamma) and (pgamma) implementations respectively, in the stats package [109, 110].

#### Model validation

As described above, the fuzzy logic approach infers a regulatory model to consist of an output node, input nodes and respectively derived regulatory rule that relate each input node to the output node. We validated derived models for each feature output in the independent *GLI1* siRNA knockdown experiments datasets generated by Falkenberg et al (2016). In this dataset, the authors focused on the genes *GLI1* and *PSMD13* as potential vorinostat-resistance candidate genes, identified from previous screens. Falkenberg and colleagues performed transcriptome analysis on vorinostat-resistant HCT116 cells (HCT116-VR) upon knockdown of these candidate genes in the presence and absence of vorinostat. According to the authors, treatment of vorinostat-resistant cells with the *GLI1* small-molecule inhibitor, GANT61, **phenocopied** the effect of *GLI1* knockdown. Therefore, for independent validation of our inferred regulatory models, we reason that for model estimated fit in the test data should as closely as possible be similar to (or better than the) estimated fit in the training dataset. The two timepoints for drug treatment assessed by Falkenberg and colleague represent a timepoint before induction of apoptosis (4hrs for siGLI1) and a timepoint when apoptosis could be detected (8hrs for siGLI1). Therefore for this validation, we used the sample expression data at 8hrs (see the table 4).

**Table 4:** Independent Validation Dataset

|  | siRNARx | drugRx | timepoint |
| --- | --- | --- | --- |
| SRX548958 | siGLI1 | vorinostat | 8hr |
| SRX548972 | siGLI1 | vorinostat | 8hr |
| SRX548986 | siGLI1 | vorinostat | 8hr |

#### Network construction and validation

For each output node, the best-fitted model as determined by estimated fit difference between the associated models in the training and validation data was selected as a representative model. Representative models were consolidated into a single regulatory network (Figure 3). We reasoned that, models with minimal estimated fit difference are more likely stable than those with high differences.

**Figure 3:**
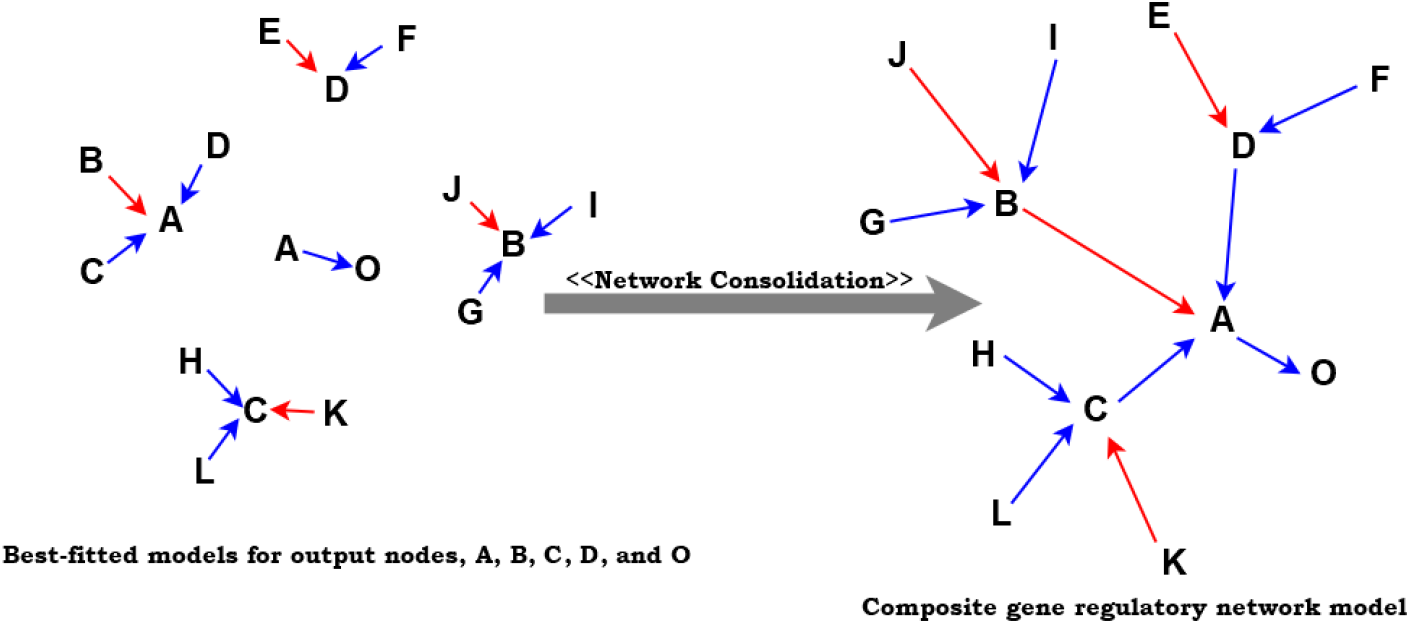
Regulatory network construction – constructed from consolidation of representative best-fitted models for all output nodes

To validate the derived regulatory network, we compared the monotonic and adaptive changes[111] observed by a dynamic simulation of the network over 5, 000 time-step iterations in the training data against that observed in the validation data. We reasoned that the distribution of observed changes between the training data network simulation and the independent validation data simulation would not be significantly different.

To simulate the network, we derived successive time-step expression values (*I*_*n*+1_) for each node by a linear combination of the previous (*I*_*n*-1_) and new values (*I_n_*), to ensure the system converge smoothly towards equilibrium[99]. Given by Gormley et al, new values (*I_n_*) were computed as:

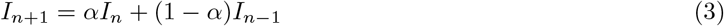

Where the *α* option specifies the ‘mixing parameter’, guiding how quuickly the simulation reaches system equilibrium. New values for each node were based on the initial conditions and the fuzzy relations (regulatory rules) inferred from the training data. Zhang et al (2019) respectively described monotonic *S_M_* and adaptive changes *S_A_* as:

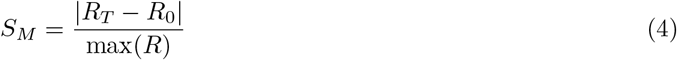

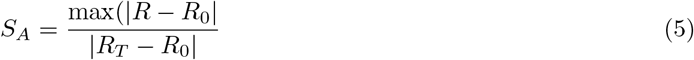

Where *R* are the estimated values over the entire iteration, *R*_0_ are observed values at the start of simulation and *R_T_* are values observed at the end of simulation. We utilized the `Student t-test` to determine if there is any difference in monotonic and adaptive network simulation changes between the training data and independent network validation data. To effectively simulate a knockdown and making the validation dataset-2 more comparable, we in-silico kept the level of knocked-down feature expression unchanged throughout the simulation steps. The table (Table 5) shows the dataset considered for independent validation of regulatory network (validation dataset-2).

**Table 5:** Independent Validation Dataset – in-silico knockout simulation

|  | siRNARx | drugRx | timepoint |
| --- | --- | --- | --- |
| SRX548952 | mock | vorinostat | 4hr |
| SRX548953 | mock | vorinostat | 8hr |
| SRX548954 | mock | vorinostat | 12hr |
| SRX548966 | mock | vorinostat | 4hr |
| SRX548967 | mock | vorinostat | 8hr |
| SRX548968 | mock | vorinostat | 12hr |
| SRX548980 | mock | vorinostat | 4hr |
| SRX548981 | mock | vorinostat | 8hr |
| SRX548982 | mock | vorinostat | 12hr |

#### Clinical Significance Evaluation

To evaluate biomedical significance of inferred regulatory network, we first estimated importance of all nodes contained therein. We defined node importance score (*I_i_*) similar to Zhang et al’s [111]. The node importance score estimates integrate network topology, network edge interaction strengths and gene expression. To encapsulate these, Zhang and colleagues defined a hub score (*H*), a local network entropy (*S*) and an adaptation score (*A*) and integrated these into a comprehensive index for each node – a normalized rank sum of these values.

A Hub score assesses a node’s connectivity to other nodes. It is the principal eigenvector of the adjacency matrix of the inferred regulatory network. If

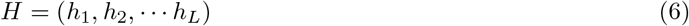

Zhang et al described the hub score of node *i* as *h_i_*.

Extending the works of Teschendorff and Severini [112], Zhang et al described local entropies as the degree of randomness in the local pattern of information flux around each node[111]. This is analogous to the centrality entropy described by Ortiz-Arroyo and Hussein [113]. It is a measure of the centrality of nodes depending on their contribution to the entropy of the derived regulatory network. We computed each nodes local entropy using Jalili et al’ scentiserve R package implementation of entropy[114]; derived from Shannon’s [115] definition of entropy which states that the entropy of a random variable *X* that can take *n* values is:

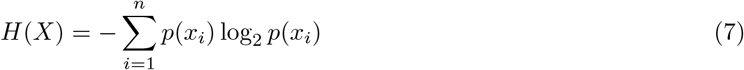

Jalili et al’s centrality entropy measure *H_ce_* of a graph G, is defined as:

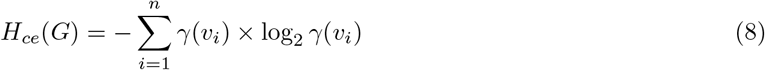

where 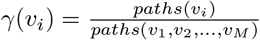 where *paths*(*v_i_*) is the number of geodesic paths from node *v_i_* to all the other nodes in the graph and *paths*(*v*_1_, *v*_2_,…, *v_M_*) is the total number of geodesic paths M that exists across all the nodes in the graph.

In place of an adaptation score rank, we modified the node importance score to include instead the fit rank (*r^F^*), the mean edges confidences rank (*r^E^*) and the delta rank (*r^D^*). We defined the fit rank as the rank of the estimated fit associated with the respective node in the network. We defined the mean edges confidences rank as the rank of the average of edge confidences returned from the STRING database associated with the node and contained in the node’s regulatory model inferred by the fuzzy logic approach. To moderate the estimated fits, we defined the delta rank as the rank of the difference in model-associated estimated fits observed in the training and independent validation datasets.

We defined an importance score (*I_i_*) for each node as the normalized rank sum of these values, similar to Zhang et al’s.

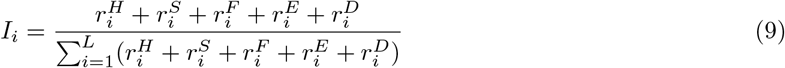

Similar to Zhang and colleagues’[111], we evaluated the potential for highly ranked regulatory node features or themes to predict short-(three or less years) and mid-term survival (greater than 3 years). We reasoned that these features are potentially able to drive tumor cells to either circumvent or succumb to epistatic events. We fitted a logistic regression model using the expression profile and clinical information we retrieved on the cancer genome atlas (TCGA) primary colorectal cancer samples – incorporating our derived node importance measures as penalty weights and specifying the 3-year survival statuses (dead or alive) as the outcome. Given *y_i_* = 0 or 1 as the binary response outcome associated with the *i*-th sample in *n* patients; *p_i_* = Pr(*y_i_* = 1); *i* = 1, ···, *n*; and *x_i_* = (*x*_*i*1_, *x*_*i*2_,…, *x_iL_*)^*T*^ is the expression profiles of the genes in the *i*-th patient, we modeled the logistic regression model as:

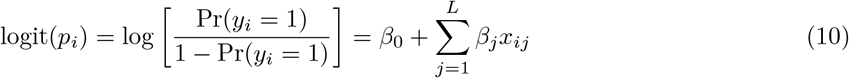

where *β*_0_ and *β_j_* are respectively the intercept and regression coefficients.

We randomly divided the data into training and test subdatasets at varying sample ratios of 50%, 60%, 70%, and 80%. We ran 100 repeated estimates at the different sample ratios. We calculated the areas under the ROC curves (AUCs) for the training and test dataset. We further evaluted the association of the top ranked features with survival using a Kaplan–Meier (K-M) survival analysis[116–118] and estimated significance between the K-M curves using the Cox proportional hazard model[119] and the two-sided log-rank test[120]. We classified patients into two groups (high-risk vs low-risk) based on the optimal cutoff using the ROC approach.

## Results

### RNA Sequence Analyses

From a total of 45 samples in the GSE56788 dataset, 34 samples succesfully passed through our analysis pipeline. 11 samples failed because of potentially corrupted raw data files. Samples that passed are shown in Table 6. These include two assays each of the interferring RNA treatment samples siCCNK, siEIF3L, siGLI1, siJAK2, siNFYA, siPOLR2D, siPSMD13 and siRGS18; one assay of the siBEGAIN treatment sample; and, three assays each for the interferring RNA treatment samples siCDK10, siDPPA5, siSAP130, siTGM5 and siTOX4. Two assays were experiment control samples. Spannig gene products involved in the cell cycle-, gene transcription- and signal transduction pathways-associated biological processes, Table 7 shows the siRNA targeted (knocked-down) genes in the vorinotat-resistant colon cancer cell line sythetic lethality experiment assays.

**Table 6:** GSE56788 dataset QC assessment samples

|  | Sample | Treatment |
| --- | --- | --- |
| 1 | SRX516756.sra_data | siCCNK |
| 2 | SRX516757.sra_data | siCDK10 |
| 3 | SRX516758.sra_data | siDPPA5 |
| 4 | SRX516759.sra_data | siEIF3L |
| 5 | SRX516760.sra_data | siGLI1 |
| 6 | SRX516761.sra_data | siJAK2 |
| 7 | SRX516762.sra_data | siNFYA |
| 8 | SRX516763.sra_data | siPOLR2D |
| 9 | SRX516764.sra_data | siPSMD13 |
| 10 | SRX516765.sra_data | siRGS18 |
| 11 | SRX516766.sra_data | siSAP130 |
| 12 | SRX516767.sra_data | siTGM5 |
| 13 | SRX516768.sra_data | siTOX4 |
| 14 | SRX516769.sra_data | mock |
| 15 | SRX516772.sra_data | siCDK10 |
| 16 | SRX516773.sra_data | siDPPA5 |
| 17 | SRX516781.sra_data | siSAP130 |
| 18 | SRX516782.sra_data | siTGM5 |
| 19 | SRX516783.sra_data | siTOX4 |
| 20 | SRX516784.sra_data | mock |
| 21 | SRX516785.sra_data | siBEGAIN |
| 22 | SRX516786.sra_data | siCCNK |
| 23 | SRX516787.sra_data | siCDK10 |
| 24 | SRX516788.sra_data | siDPPA5 |
| 25 | SRX516789.sra_data | siEIF3L |
| 26 | SRX516790.sra_data | siGLI1 |
| 27 | SRX516791.sra_data | siJAK2 |
| 28 | SRX516792.sra_data | siNFYA |
| 29 | SRX516793.sra_data | siPOLR2D |
| 30 | SRX516794.sra_data | siPSMD13 |
| 31 | SRX516795.sra_data | siRGS18 |
| 32 | SRX516796.sra_data | siSAP130 |
| 33 | SRX516797.sra_data | siTGM5 |
| 34 | SRX516798.sra_data | siTOX4 |

**Table 7:** siRNA Experiments Targetted Genes

|  | SYMBOL | GENENAME | ENTREZID |
| --- | --- | --- | --- |
| 1 | BEGAIN | brain enriched guanylate kinase associated | 57596 |
| 2 | CCNK | cyclin K | 8812 |
| 3 | CDK10 | cyclin dependent kinase 10 | 8558 |
| 4 | DPPA5 | developmental pluripotency associated 5 | 340168 |
| 5 | EIF3L | eukaryotic translation initiation factor 3 subunit L | 51386 |
| 6 | GLI1 | GLI family zinc finger 1 | 2735 |
| 7 | JAK2 | Janus kinase 2 | 3717 |
| 8 | NFYA | nuclear transcription factor Y subunit alpha | 4800 |
| 9 | POLR2D | RNA polymerase II subunit D | 5433 |
| 10 | PSMD13 | proteasome 26S subunit, non-ATPase 13 | 5719 |
| 11 | RGS18 | regulator of G protein signaling 18 | 64407 |
| 12 | SAP130 | Sin3A associated protein 130 | 79595 |
| 13 | TGM5 | transglutaminase 5 | 9333 |
| 14 | TOX4 | TOX high mobility group box family member 4 | 9878 |

#### Reads quality assessment

All quality assessment measures as defined by Andrews and colleagues at the Babraham Institute[61], except the ‘Per base sequence content’ module which represents the relative amount of each base in the entire genome (Table 8), were satisfied. These included the: ‘Adapter Content’, ‘Overrepresented sequences’, ‘Per base N content’, ‘Per base sequence content’, ‘Per base sequence quality’, ‘Per sequence GC content’, ‘Per sequence quality scores’, ‘Sequence Duplication Levels, and the ‘Sequence Length Distribution’ module passed QC assessment. In most instances, the ‘Per base sequence content’ is not of biological concern as this arises from technical issues relating to using primers with random hexamers or the use of transposases which are biased toward specific cleavage sites during the library generation step. It is understood that, because of this some biases may occur, particularly at the start of the sequence reads[61]. The ‘Per base sequence content’ failure is triggered if the difference between A and T, or G and C is greater than 20% in any position. This is generally not a problem if the biases and failures can be visualized (see Figure 4) and attributed to around the first 12 base locations.

**Figure 4:**
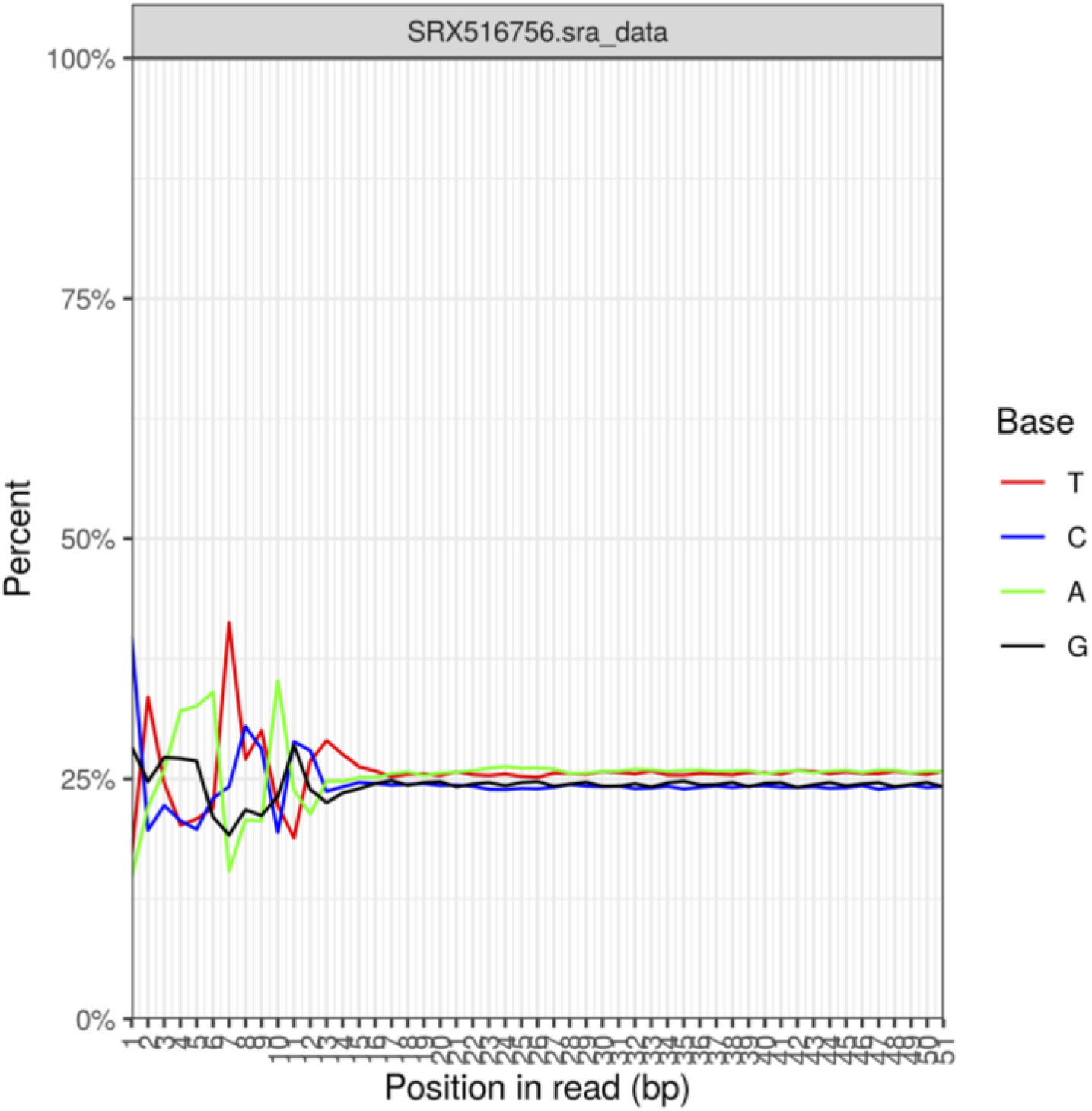
Per Base Sequence Content Plot - the proportion of each each of the four normal DNA bases at each sequence read position in the SRX516756.sra_data (siCCNK) experiment sample

**Table 8:**
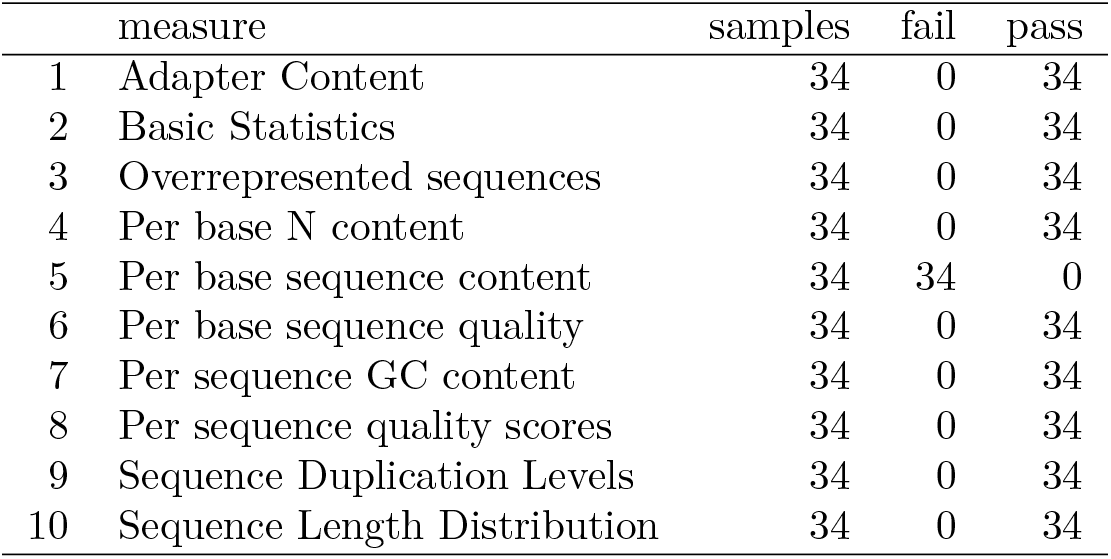
GSE56788 dataset QC assessment by quality control measures

|  | measure | samples | fail | pass |
| --- | --- | --- | --- | --- |
| 1 | Adapter Content | 34 | 0 | 34 |
| 2 | Basic Statistics | 34 | 0 | 34 |
| 3 | Overrepresented sequences | 34 | 0 | 34 |
| 4 | Per base N content | 34 | 0 | 34 |
| 5 | Per base sequence content | 34 | 34 | 0 |
| 6 | Per base sequence quality | 34 | 0 | 34 |
| 7 | Per sequence GC content | 34 | 0 | 34 |
| 8 | Per sequence quality scores | 34 | 0 | 34 |
| 9 | Sequence Duplication Levels | 34 | 0 | 34 |
| 10 | Sequence Length Distribution | 34 | 0 | 34 |

Figure 5 shows the Per Base Sequence Quality plot for the SRX516756.sra_data. This shows the distribution of quality scores for bases at the respective positions in a box plot with whiskers. The y-axis shows the quality scores. A better base call is indicated by a higher score. The background of the graph divides the y axis into very good quality (green), reasonable quality (orange), and poor quality (red) calls. It is not unusual for the quality of a base call to degrade toward the end of the read. Although scores appear generally okay, it can be seen that the scores reported at the first 5 —10 base positions are of lesser quality.

**Figure 5:**
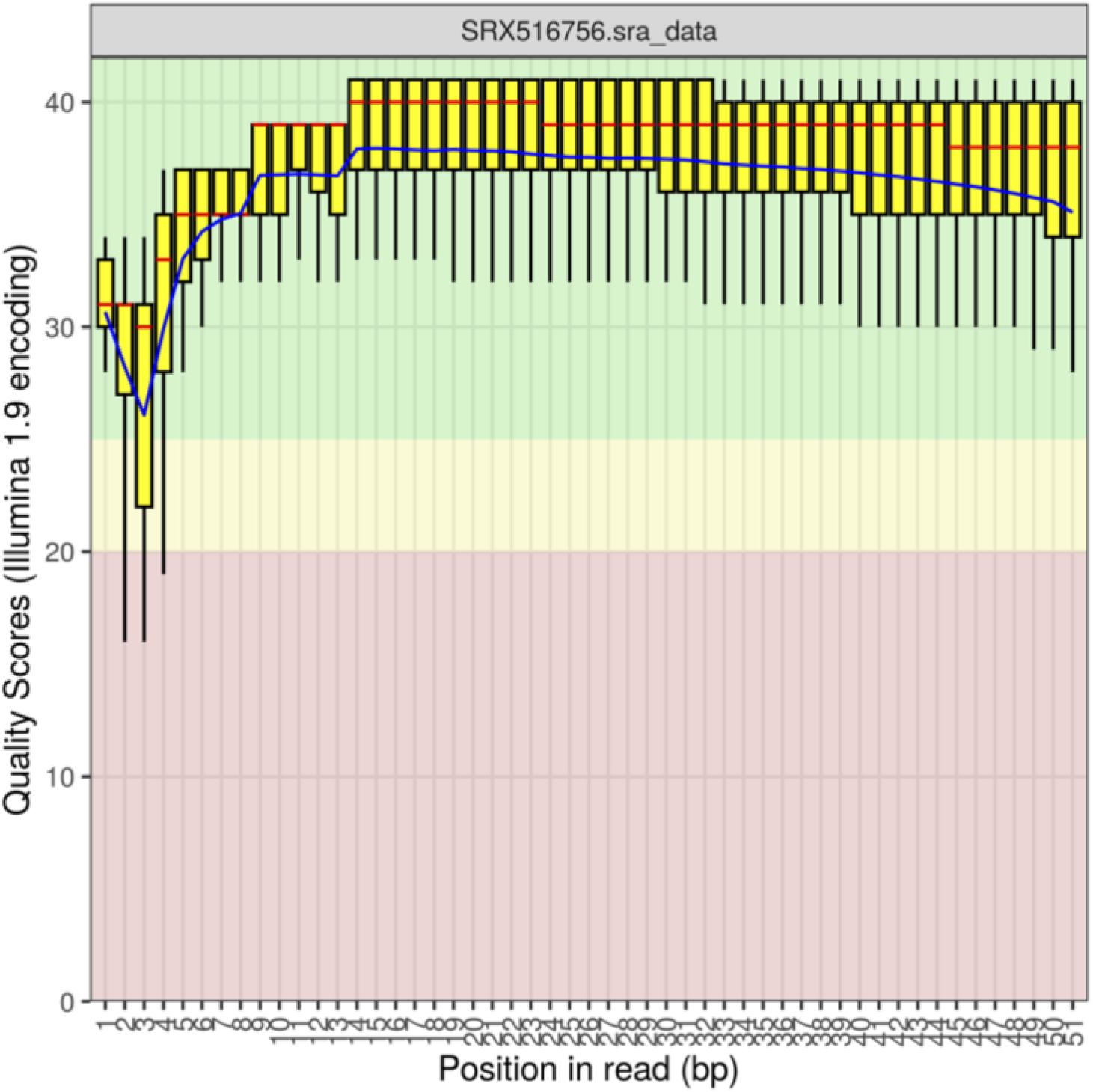
Per Base Sequence Quality Plot - the distribution of quality scores for bases at the respective positions in the SRX516756.sra_data experiment sample

Table 9 shows estimates for quality assessment parameters in each RNA sequencing sample in the GSE56788 dataset. Assessed parameters included percent duplication, percent GC content, and average sequence read length. Average duplication rate is estimated to be 20.21%, while that of GC content stood at 48.68%.

**Table 9:** GSE56788 dataset QC assessment of sequence reads

|  | Sample | Duplicates(%) | GC% | Length |
| --- | --- | --- | --- | --- |
| 1 | SRX516756.sra_data | 23.64 | 48.00 | 51 |
| 2 | SRX516757.sra_data | 17.62 | 48.00 | 51 |
| 3 | SRX516758.sra_data | 18.78 | 49.00 | 51 |
| 4 | SRX516759.sra_data | 19.32 | 48.00 | 51 |
| 5 | SRX516760.sra_data | 22.75 | 48.00 | 51 |
| 6 | SRX516761.sra_data | 21.38 | 49.00 | 51 |
| 7 | SRX516762.sra_data | 17.91 | 49.00 | 51 |
| 8 | SRX516763.sra_data | 23.47 | 49.00 | 51 |
| 9 | SRX516764.sra_data | 18.53 | 48.00 | 51 |
| 10 | SRX516765.sra_data | 20.36 | 49.00 | 51 |
| 11 | SRX516766.sra_data | 19.52 | 49.00 | 51 |
| 12 | SRX516767.sra_data | 18.08 | 49.00 | 51 |
| 13 | SRX516768.sra_data | 20.73 | 48.00 | 51 |
| 14 | SRX516769.sra_data | 18.67 | 49.00 | 51 |
| 15 | SRX516772.sra_data | 18.30 | 48.00 | 51 |
| 16 | SRX516773.sra_data | 18.26 | 49.00 | 51 |
| 17 | SRX516781.sra_data | 20.91 | 49.00 | 51 |
| 18 | SRX516782.sra_data | 20.16 | 49.00 | 51 |
| 19 | SRX516783.sra_data | 19.73 | 49.00 | 51 |
| 20 | SRX516784.sra_data | 17.66 | 49.00 | 51 |
| 21 | SRX516785.sra_data | 24.11 | 49.00 | 51 |
| 22 | SRX516786.sra_data | 23.17 | 48.00 | 51 |
| 23 | SRX516787.sra_data | 20.90 | 49.00 | 51 |
| 24 | SRX516788.sra_data | 19.87 | 49.00 | 51 |
| 25 | SRX516789.sra_data | 19.26 | 48.00 | 51 |
| 26 | SRX516790.sra_data | 19.57 | 49.00 | 51 |
| 27 | SRX516791.sra_data | 20.42 | 49.00 | 51 |
| 28 | SRX516792.sra_data | 18.01 | 49.00 | 51 |
| 29 | SRX516793.sra_data | 24.32 | 49.00 | 51 |
| 30 | SRX516794.sra_data | 19.15 | 49.00 | 51 |
| 31 | SRX516795.sra_data | 18.77 | 48.00 | 51 |
| 32 | SRX516796.sra_data | 23.23 | 49.00 | 51 |
| 33 | SRX516797.sra_data | 17.94 | 49.00 | 51 |
| 34 | SRX516798.sra_data | 22.56 | 48.00 | 51 |

#### Reads quantification

Maximum number of reads was found to be 26430192 reads in the SRX516756.sra_data (siCCNK) sample, while the minimum was 9756110, in the SRX516790.sra_data (siGLI1) sample - an approximately three fold difference across the sythetic lethal experiment assays (Fig. ??, Table 10). As opposed to 45 samples, results are presented for 34 samples. As previously mentioned, data for 11 samples failed on `topHat2` alignment on execution, potentially due to corrupted samples’ raw data file.

**Table 10:** GSE56788 dataset sequence reads

|  | Sample | Treatment | Total |
| --- | --- | --- | --- |
| 1 | SRX516756.sra_data | siCCNK | 26430192 |
| 2 | SRX516757.sra_data | siCDK10 | 11926761 |
| 3 | SRX516758.sra_data | siDPPA5 | 14012687 |
| 4 | SRX516759.sra_data | siEIF3L | 17264395 |
| 5 | SRX516760.sra_data | siGLI1 | 16841563 |
| 6 | SRX516761.sra_data | siJAK2 | 19008717 |
| 7 | SRX516762.sra_data | siNFYA | 12986269 |
| 8 | SRX516763.sra_data | siPOLR2D | 16712374 |
| 9 | SRX516764.sra_data | siPSMD13 | 13782737 |
| 10 | SRX516765.sra_data | siRGS18 | 16399848 |
| 11 | SRX516766.sra_data | siSAP130 | 14471644 |
| 12 | SRX516767.sra_data | siTGM5 | 14561182 |
| 13 | SRX516768.sra_data | siTOX4 | 17051622 |
| 14 | SRX516769.sra_data | mock | 14581166 |
| 15 | SRX516772.sra_data | siCDK10 | 12726019 |
| 16 | SRX516773.sra_data | siDPPA5 | 12870208 |
| 17 | SRX516781.sra_data | siSAP130 | 14235470 |
| 18 | SRX516782.sra_data | siTGM5 | 13362627 |
| 19 | SRX516783.sra_data | siTOX4 | 13970706 |
| 20 | SRX516784.sra_data | mock | 11168125 |
| 21 | SRX516785.sra_data | siBEGAIN | 23372015 |
| 22 | SRX516786.sra_data | siCCNK | 17367956 |
| 23 | SRX516787.sra_data | siCDK10 | 15621563 |
| 24 | SRX516788.sra_data | siDPPA5 | 14589712 |
| 25 | SRX516789.sra_data | siEIF3L | 14106493 |
| 26 | SRX516790.sra_data | siGLI1 | 9756110 |
| 27 | SRX516791.sra_data | siJAK2 | 15299261 |
| 28 | SRX516792.sra_data | siNFYA | 11219308 |
| 29 | SRX516793.sra_data | siPOLR2D | 15782493 |
| 30 | SRX516794.sra_data | siPSMD13 | 12804888 |
| 31 | SRX516795.sra_data | siRGS18 | 12792707 |
| 32 | SRX516796.sra_data | siSAP130 | 18050180 |
| 33 | SRX516797.sra_data | siTGM5 | 12923634 |
| 34 | SRX516798.sra_data | siTOX4 | 17513707 |

Similar QC assessment profile is observed for the data in both the GSE56788 and the GSE57871 datasets.\

### Feature Selection, for Regulatory Network Inference

#### Features’ mean absolute deviations (MADs)

As previously described, we reasoned that the most changing features, in terms of expression values across pertubations, are more informative in the context of regulatory networks than non-changing features (see section). To determine these features, we estimated the mean abosolute deviations of each of the 28, 395 genes across all available assay samples in our model-inferring (training) dataset. The observed median of MADs is 0.2033 (Mean=0.2905, SD=0.3482). With a maximum and minimum observed MAD of 2.1333 and 0.0 respectively, over 30% (11, 123) of features do not appear to change (Fig 6, 1st Qu=0.0000). These include those for knocked-down features, DPPA5, RGS18, and TGM5. The feature with maximum MAD was HRNR (hornerin).

**Figure 6:**
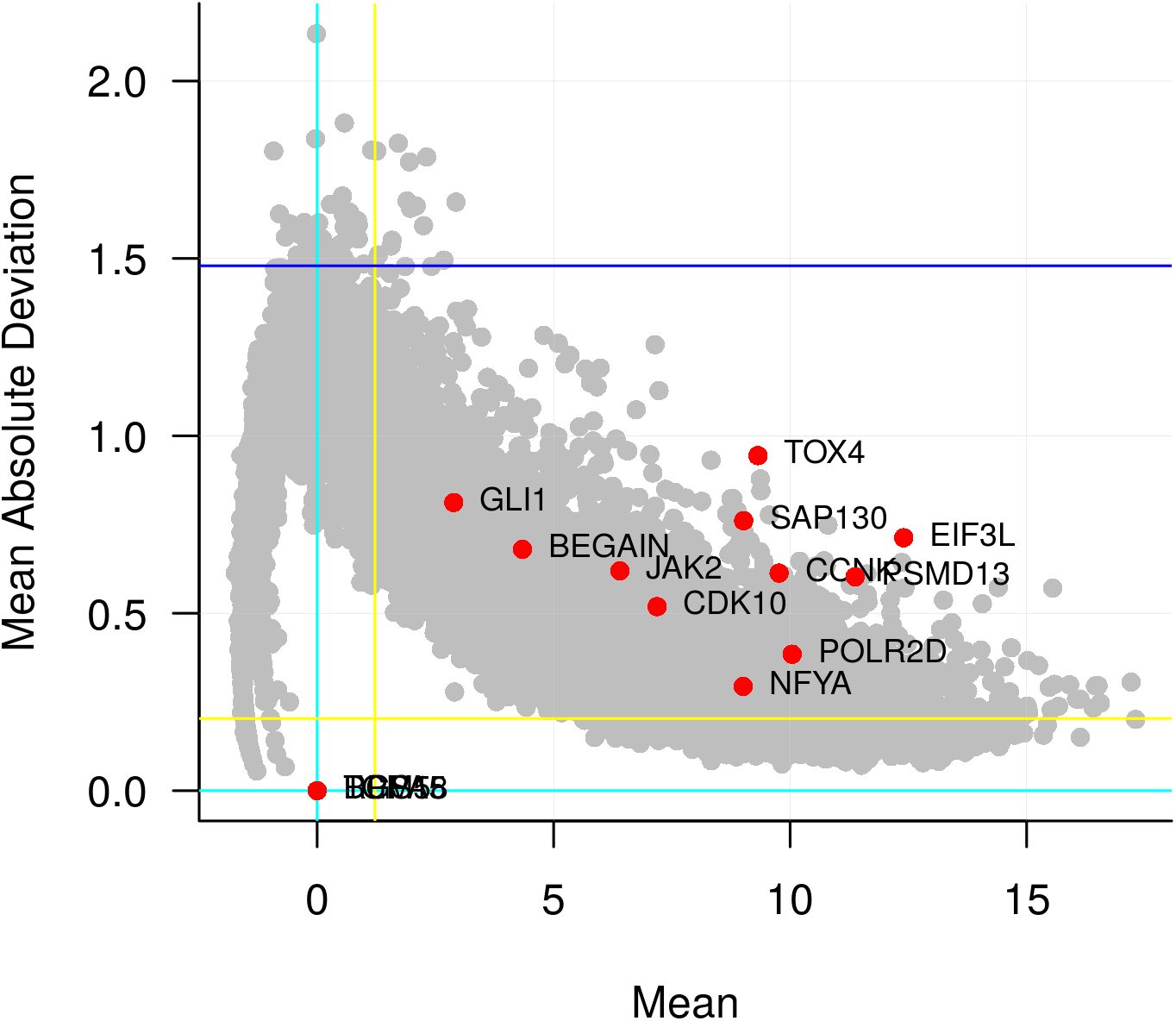
A plot of mean absolute deviations versus mean of log expresson values across vorinostat-resistant colon cancer synthetic lethal experiment assays. Coordinates of siRNA-targeted (knocked-down) features in the different assays are indicated in red. The yellow lines indicate the median of the estimated mean absolute deviations and the median of the mean log-expressions respectively.

#### Features’ differential expression

Another measure of change we employed was the differential expression for each feature – an estimate of features that are truly different in terms of expression values between conditions. We estimated the differential expression of features in each knock-down assay group against the control assays. At adjusted p-values ≤0.05, the maximum number of differentially expressed features (8, 055) were found in the POLR2D siRNA knockdown assays, while the least number (2, 504) were found in the RGS18 knockdown experiments (Table 11, Fig. 7 and 8). 1, 645 features were differentially expressed in 5 comparisons of siRNA knockdown assays versus control assays, while 6 features are differentially expressed in all comparisons(Fig. 9). The features found to be differentially expressed in all contrasts include the NRBP1, MTHFD2, ALDH1A3, TNS2, LIMA1 and the BAK1 gene products. Cummulatively 13, 090 features were found to be differentially expressed in at least one contrast comparison, while 4, 270 were found to be differentially expressed in at least half of the comparisons (Fig. 10, Table 12). In terms of features discovered to be differentially expressed, the assay groups appear to cluster into 3 major groups (Fig. 11).

**Figure 7:**
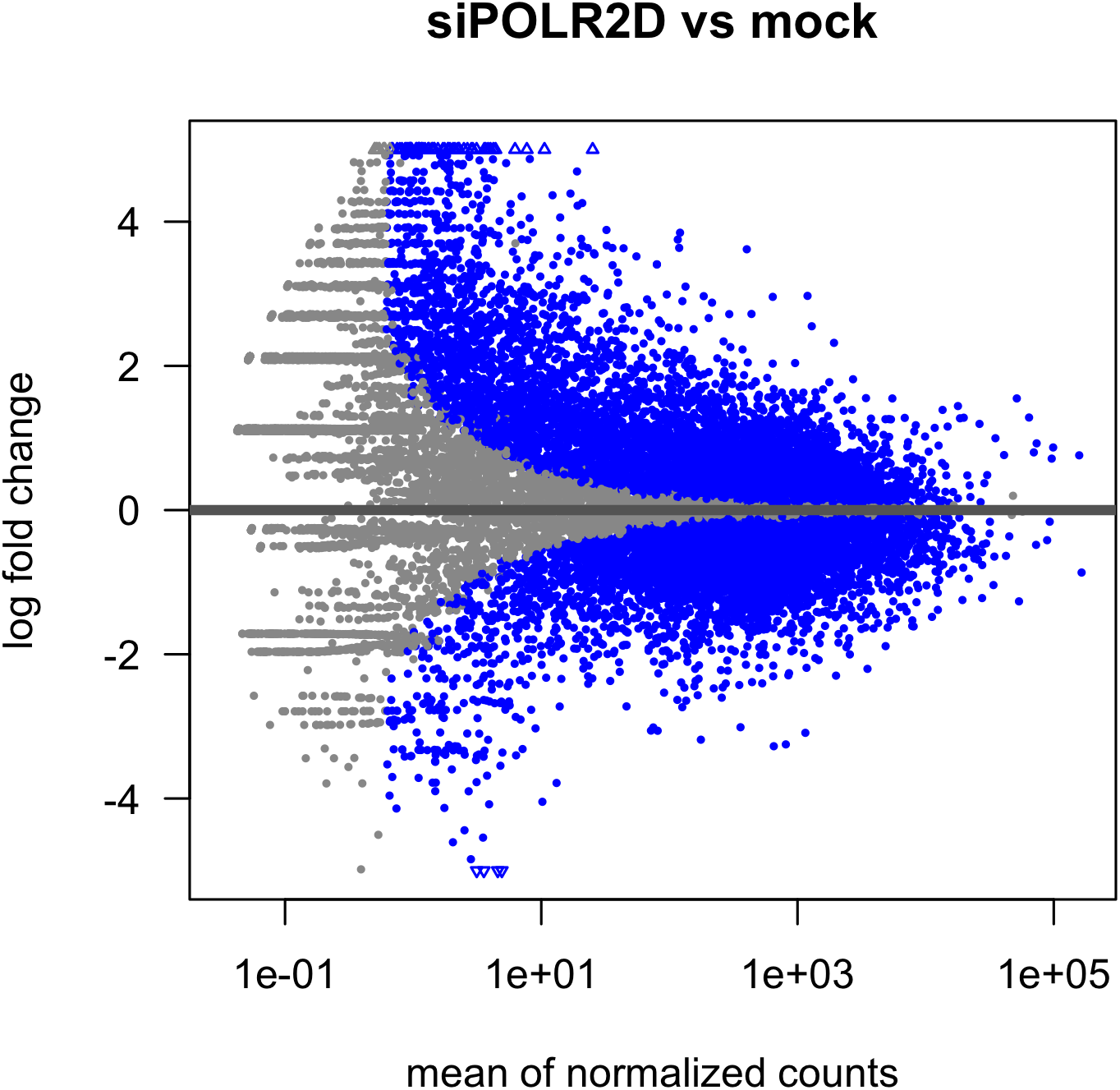
MA Plot highligting computed differentially expressed features (blue) between the POLR2D siRNA knockdown assays and the control (mock siRNA) assays

**Figure 8:**
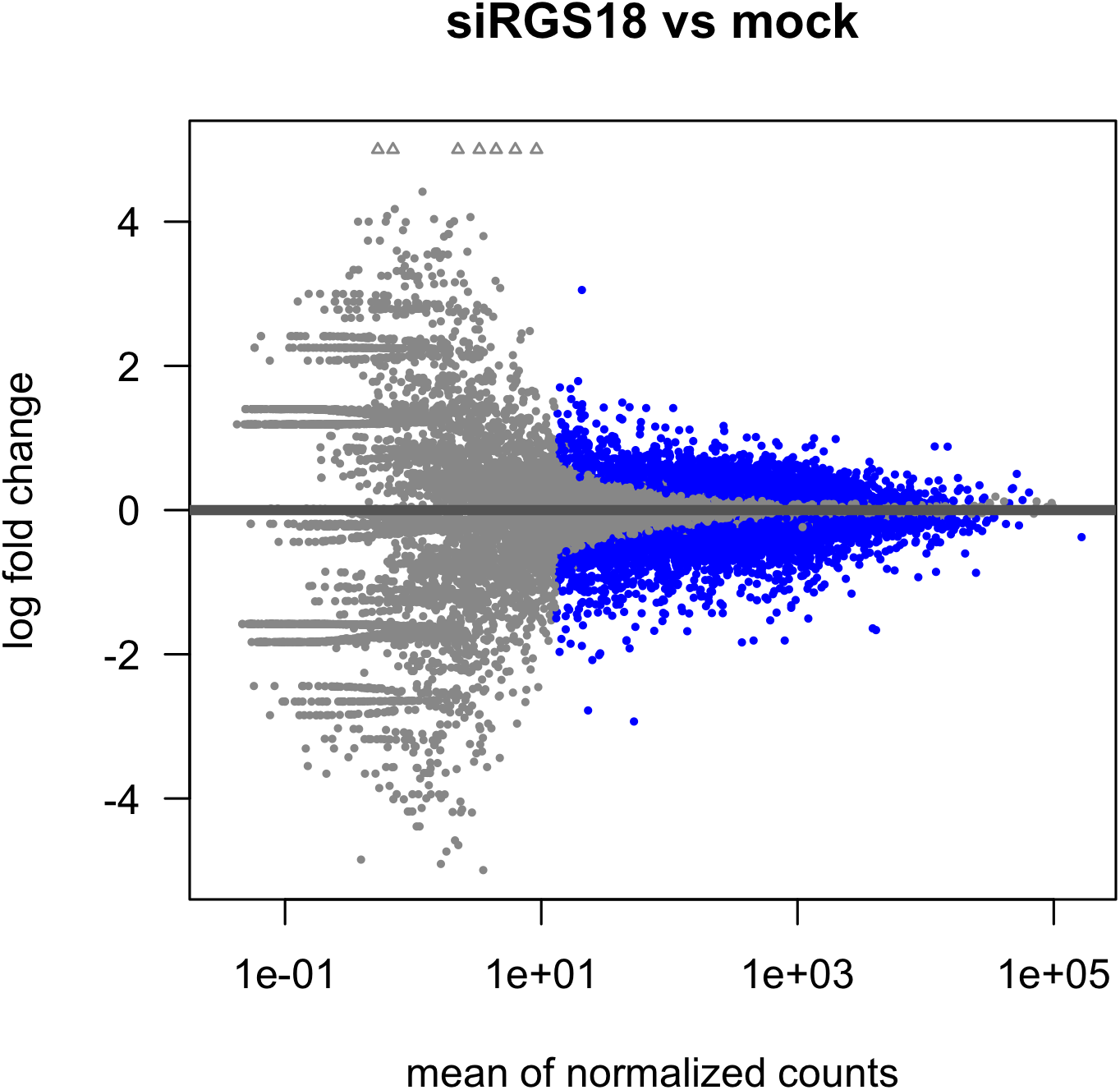
MA Plot highligting computed differentially expressed features (blue) between the RGS18 siRNA knockdown assays and the control (mock siRNA) assays

**Figure 9:**
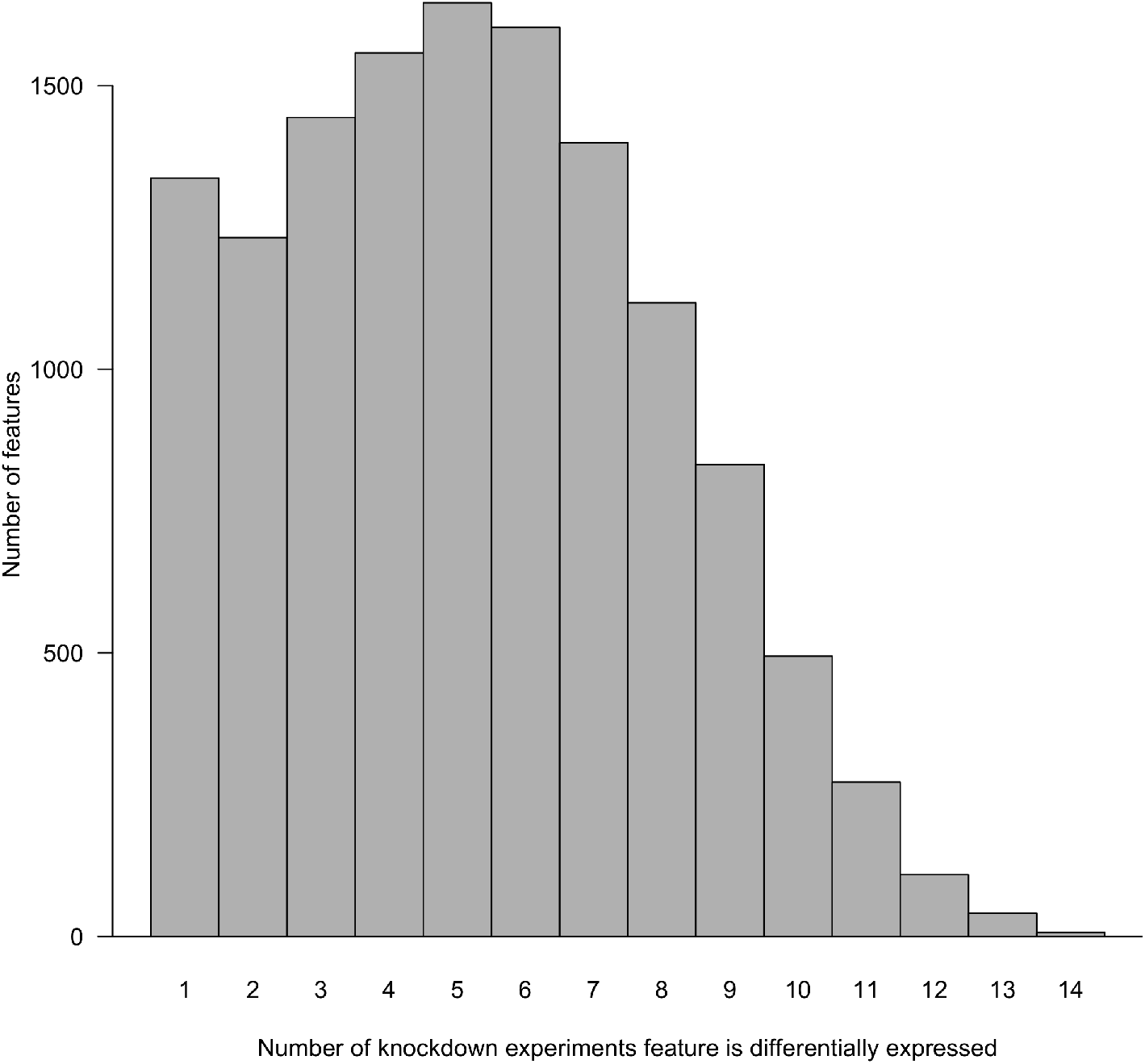
A barplot showing the number of features found to be differentially expressed in the comparisons against the control experiments. 6 features are found to be differentially expressed in all 14 comparisons of siRNA knockdown experiments versus the controls. Most of the differentially expressed features are found in a least 5 comparisons.

**Figure 10:**
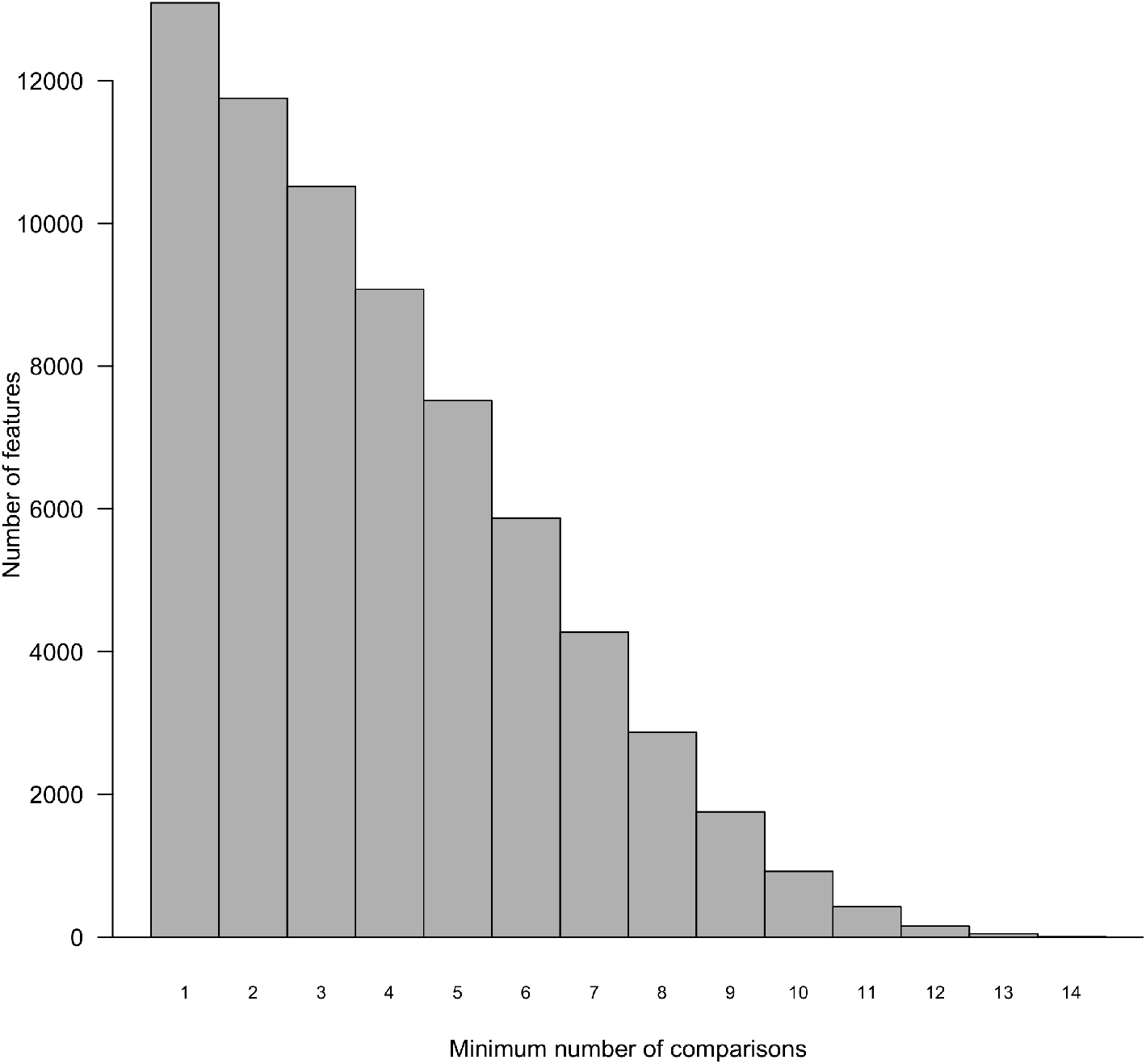
Barplot showing the cummulative number of features found to be differentially expressed in the comparisons against the control experiments. 6 features are found to be differentially expressed in all 14 comparisons of siRNA knockdown experiments versus the controls. Cummulatively 13, 090 features were found to be differentially expressed in at least one comparison, while 4, 270 were found to be differentially expressed in at least half of the comparisons

**Figure 11:**
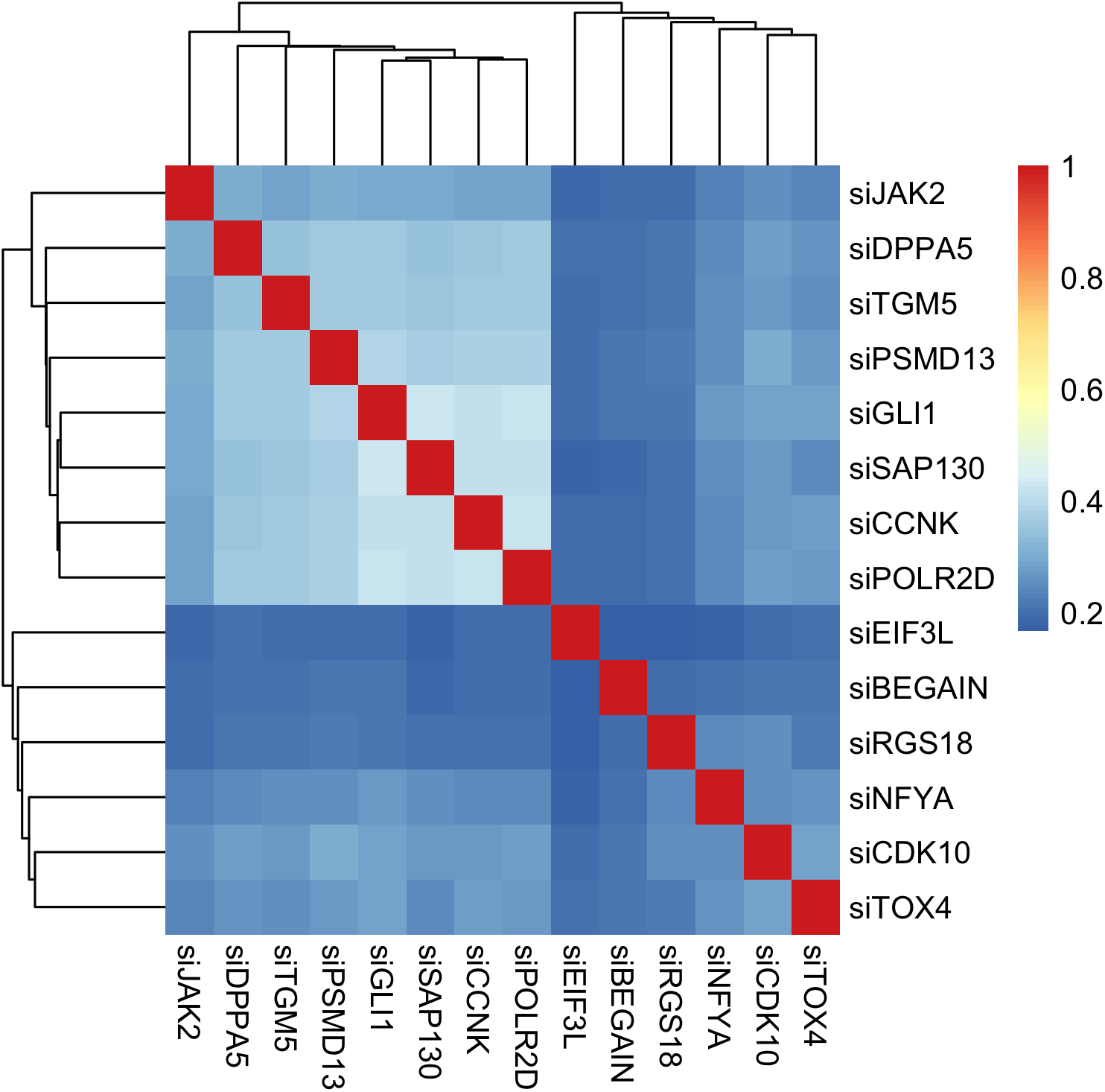
A heatmap of similarities between the siRNA knockdown assay groups, in terms of differentially expressed features. The Jaccard score estimate was used as a measure of similarities. The higher the estimated value, the more similar the groups are

**Table 11:** Number of differentially expressed features between siRNA knockdown assays and control assays

| | Differentially Expressed Features | At $\leq 0.05$ Adjusted P-value |
| --- | --- | --- |
| siCCNK | 12237 | 7068 |
| siCDK10 | 10404 | 3644 |
| siDPPA5 | 11838 | 5476 |
| siEIF3L | 10108 | 2553 |
| siGLI1 | 12409 | 6842 |
| siJAK2 | 11192 | 4035 |
| siNFYA | 10393 | 3236 |
| siPOLR2D | 12032 | 8055 |
| siPSMD13 | 11848 | 5883 |
| siRGS18 | 10456 | 2504 |
| siSAP130 | 12312 | 7540 |
| siTGM5 | 11478 | 5513 |
| siTOX4 | 11121 | 3566 |
| siBEGAIN | 10878 | 2528 |

**Table 12:** Table of cummulative occurrence of features as differentially expressed. Cummulatively, 13, 090 features are differentially expressed in at least one comparison while 6 features are differentially expressed in all 14 comparisons between the different knockdown assays versus the control experiment assays

| Comparisons | Features |
| --- | --- |
| 1 | 13090 |
| 2 | 11753 |
| 3 | 10521 |
| 4 | 9077 |
| 5 | 7519 |
| 6 | 5873 |
| 7 | 4270 |
| 8 | 2871 |
| 9 | 1754 |
| 10 | 922 |
| 11 | 428 |
| 12 | 156 |
| 13 | 47 |
| 14 | 6 |

#### Features’ expression ranges and log-fold changes

Still on evaluating features’ expression changes across knockdown experiments, we considered the log-fold change between the minimum and the maximum expression value for each feature across all knockdown assays. A large proportion of features show no log-fold change (Fig. 12). The maximum log fold change (10.436) is found in the SPP1 feature expression profile. Across all features, median log-fold change was 1.099 while mean log-fold change was 1.497. 8, 037 features have log-fold changes ≥ 2 while only 19 features have log-fold change ≥ 8 (Fig. 13 and Table 13).

**Figure 12:**
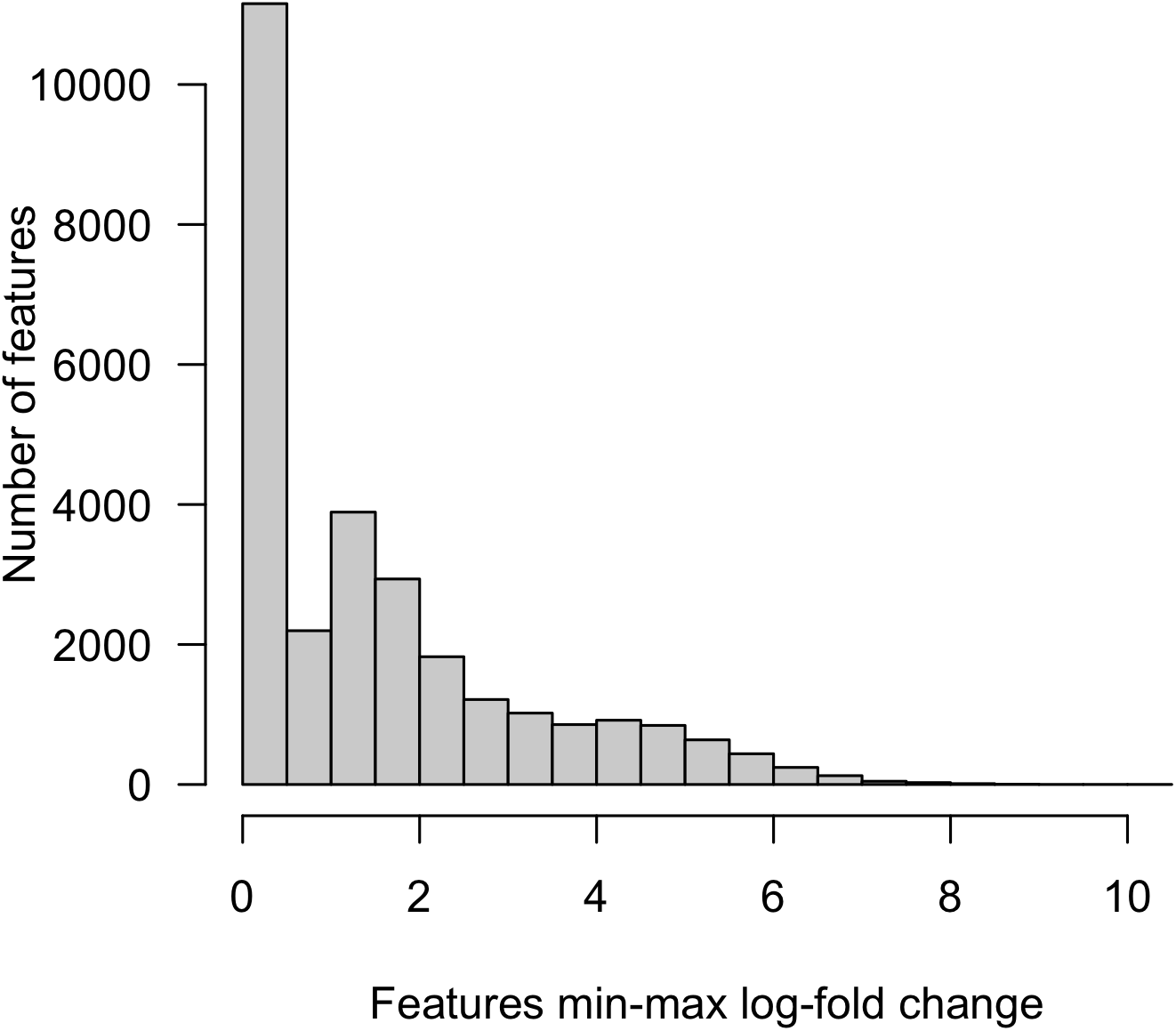
A histogram of log-fold changes between the minimum and maximum expression values for features across knockdown experiments.

**Figure 13:**
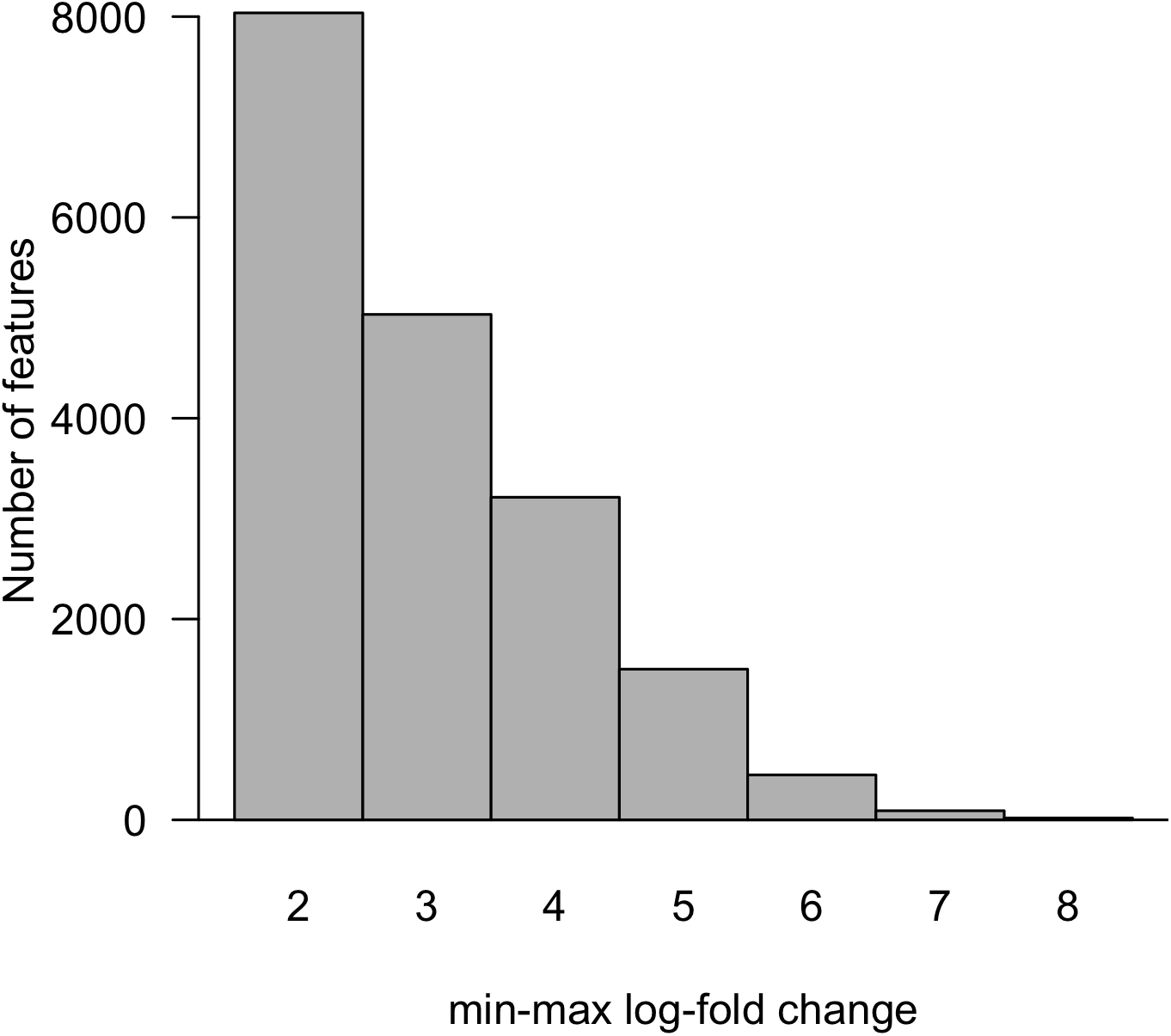
A boxplot showing the number of features with greater than or equal a value of the specified log-fold change (difference) between the minimum and maximum expression values across experiments.

**Table 13:** Number of features with min-max log-fold change with greater than or equal the specified values.

| min-max Log-fold | number of features |
| --- | --- |
| 2 | 8037 |
| 3 | 5034 |
| 4 | 3213 |
| 5 | 1501 |
| 6 | 448 |
| 7 | 91 |
| 8 | 19 |

#### Online Mendelian inheritance in man (OMIM) database features

For a more encompassing regulatory network inference feature set, and addressing limitations of purely data-driven approaches as previously mentioned, 52 features were retrieved from the online mendelian inheritance in man (OMIM) database (Table 3)[49, 50].

#### Search tool for the retrieval of interacting proteins (STRING) database search

According to criteria specified previously, and together with features determined from the OMIM database, 571 were considered to be temporally changing and potentially informative for regulatory network construction.

These were searched against the STRING database for any remote biological evidence of potential interactions – serving as a priori knowledge guide for our downstream fuzzy logic regulatory network inference. At a false discovery rate of 0.05 and minimum interation confidence of 0.150 (low confidence), retrieved interaction network consisted of 559 nodes and 8, 819 edges. Average node degree was 31.6 and the average local clustering coefficient was 0.312. With an expected number of edges of 7, 659, p-value of protein-protein interaction (PPI) enrichment was < 1e-16. With 238 mapped interactions, AKT was reported to have the most identitfied interactions. Features with only one SRING database-identified interaction were C1orf35, CCDC171, MROH8, OR51B2, PRB3, and RNF223.

#### Selected features for fuzzy logic based regulatory network

Of the 559 features found to belong to probable biological network in the STRING database, 535 were subjected to the fuzzy logic regulatory inference approach.

### Regulatory Network Inference

An inferred fuzzy logic model consists of an output node, its regulatory input nodes, and respective regulary rule that describes the interaction and relationship of the input to the output node. A regulatory rule is one of 27 rules, each represented by a three-member array notation (or tuple). The indices of the array represent respectively a low, medium and high presupposed state of the input node. Actual value (1, 2, or 3) of each element in the array represents the respective expected state (low, medium, or high) of the output node. For example, the rule [3, 2, 1] states that: when the input expression value is ‘low’, the output node is ‘high’(3); when the input is ‘medium’, the output is ‘medium’(2); and when the input is ‘high’, the output is ‘low’(1). This represents a classic repression-like regulation (i.e. negative control). The converse is true for a rule [1, 2, 3] representation. It implies that when the input expression value is ‘low’, the output node is ‘low’(1); when the input is medium, the output is medium(2); and when the input is ‘high’, the output is ‘high’(1).

This represents a classic activation-like regulation (i.e. positive control).

#### Fuzzy logic-based regulatory models

Filtering at estimated fit of 0.70, 299 output nodes and fuzzy logic regulatory models were obtained. These consist of 402 gene features. Ranked by models’ minimum difference between estimated fit in training data and independent validation data, the top models include the output nodes: TGFBR2 (model fit = 0.7011; adjusted p-value = 5.720849e-04), RIMBP3B (model fit = 0.7972; adjusted p-value = 1.907588e-05), PPP2R1A (model fit = 0.7162; adjusted p-value = 9.497439e-05), UBQLN2 (model fit = 0.7770; adjusted p-value = 4.702329e-05), WNT3A (model fit = 0.7300; adjusted p-value = 1.872238e-04), TP53 (model fit = 0.7144; adjusted p-value = 2.021138e-03), and PIK3CA (model fit = 0.7040; adjusted p-value = 7.431795e-04) amongst many others (Tables 14, 15, and 16). Enumerated to regulate the TGFBR2 gene were three regulatory inputs. These include, a inhibitory interaction (fuzzy logic rule, [2, 1, 1]) by the TNS1 gene, a stimulatory interaction (fuzzy logic rule, [1, 1, 3]) by the AXIN2 gene, and another inhibitory interaction (fuzzy logic rule, [3, 2, 2]) by the CCND1 gene products respectively. For the TP53 gene, two stimulatory interactions by the TP53I3 and the WNT1 gene products were identified (fuzzy logic rules, [1, 2, 3], and [1, 1, 3] respectively) (Tables 14, 15, and 16).

**Table 14:**
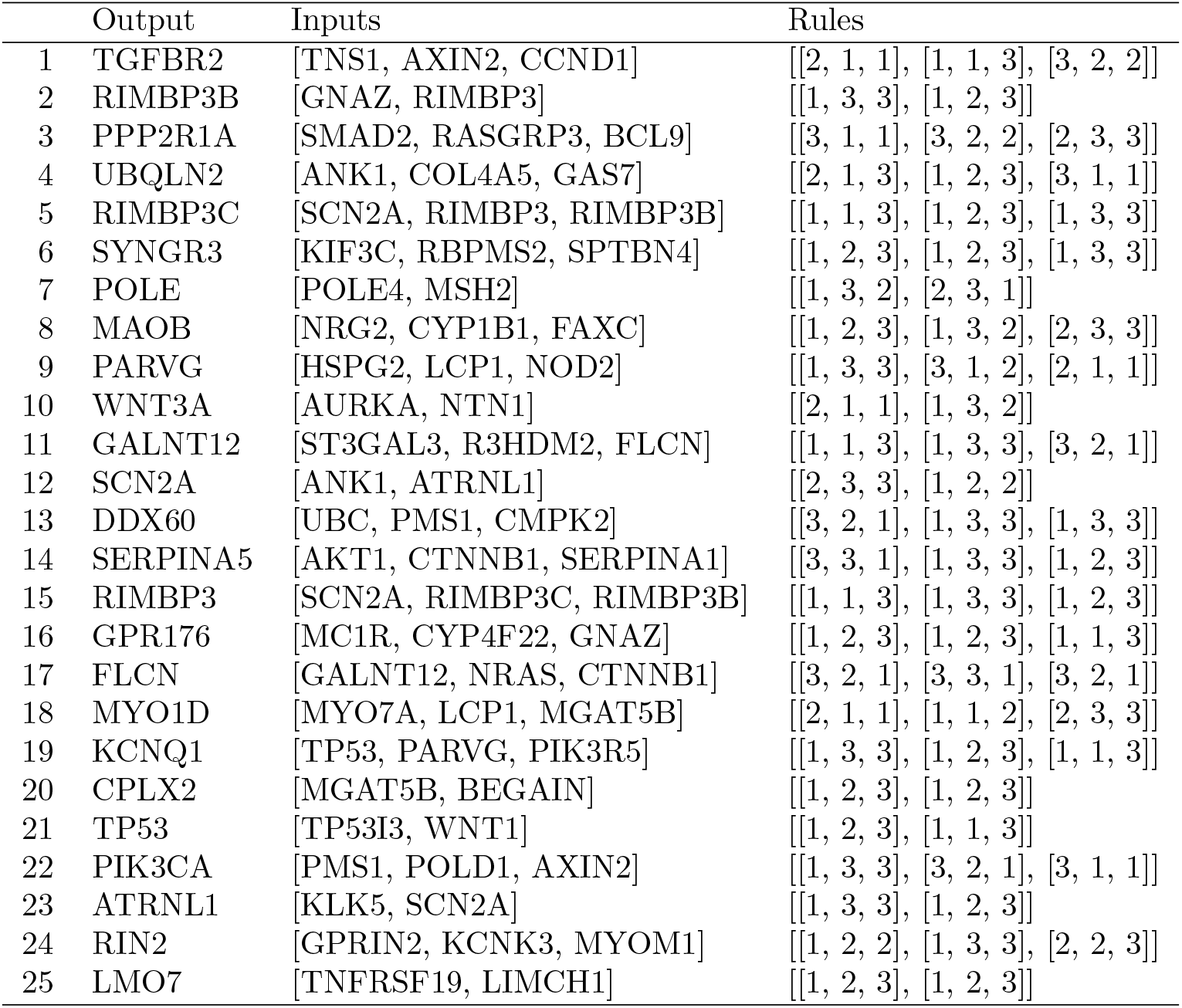
Top 25 fuzzy-logic regulatory models identified – I

|  | Output | Inputs | Rules |
| --- | --- | --- | --- |
| 1 | TGFBR2 | [TNS1, AXIN2, CCND1] | [[2, 1, 1], [1, 1, 3], [3, 2, 2]] |
| 2 | RIMBP3B | [GNAZ, RIMBP3] | [[1, 3, 3], [1, 2, 3]] |
| 3 | PPP2R1A | [SMAD2, RASGRP3, BCL9] | [[3, 1, 1], [3, 2, 2], [2, 3, 3]] |
| 4 | UBQLN2 | [ANK1, COL4A5, GAS7] | [[2, 1, 3], [1, 2, 3], [3, 1, 1]] |
| 5 | RIMBP3C | [SCN2A, RIMBP3, RIMBP3B] | [[1, 1, 3], [1, 2, 3], [1, 3, 3]] |
| 6 | SYNGR3 | [KIF3C, RBPMS2, SPTBN4] | [[1, 2, 3], [1, 2, 3], [1, 3, 3]] |
| 7 | POLE | [POLE4, MSH2] | [[1, 3, 2], [2, 3, 1]] |
| 8 | MAOB | [NRG2, CYP1B1, FAXC] | [[1, 2, 3], [1, 3, 2], [2, 3, 3]] |
| 9 | PARVG | [HSPG2, LCP1, NOD2] | [[1, 3, 3], [3, 1, 2], [2, 1, 1]] |
| 10 | WNT3A | [AURKA, NTN1] | [[2, 1, 1], [1, 3, 2]] |
| 11 | GALNT12 | [ST3GAL3, R3HDM2, FLCN] | [[1, 1, 3], [1, 3, 3], [3, 2, 1]] |
| 12 | SCN2A | [ANK1, ATRNL1] | [[2, 3, 3], [1, 2, 2]] |
| 13 | DDX60 | [UBC, PMS1, CMPK2] | [[3, 2, 1], [1, 3, 3], [1, 3, 3]] |
| 14 | SERPINA5 | [AKT1, CTNNB1, SERPINA1] | [[3, 3, 1], [1, 3, 3], [1, 2, 3]] |
| 15 | RIMBP3 | [SCN2A, RIMBP3C, RIMBP3B] | [[1, 1, 3], [1, 3, 3], [1, 2, 3]] |
| 16 | GPR176 | [MC1R, CYP4F22, GNAZ] | [[1, 2, 3], [1, 2, 3], [1, 1, 3]] |
| 17 | FLCN | [GALNT12, NRAS, CTNNB1] | [[3, 2, 1], [3, 3, 1], [3, 2, 1]] |
| 18 | MYO1D | [MYO7A, LCP1, MGAT5B] | [[2, 1, 1], [1, 1, 2], [2, 3, 3]] |
| 19 | KCNQ1 | [TP53, PARVG, PIK3R5] | [[1, 3, 3], [1, 2, 3], [1, 1, 3]] |
| 20 | CPLX2 | [MGAT5B, BEGAIN] | [[1, 2, 3], [1, 2, 3]] |
| 21 | TP53 | [TP53I3, WNT1] | [[1, 2, 3], [1, 1, 3]] |
| 22 | PIK3CA | [PMS1, POLD1, AXIN2] | [[1, 3, 3], [3, 2, 1], [3, 1, 1]] |
| 23 | ATRNL1 | [KLK5, SCN2A] | [[1, 3, 3], [1, 2, 3]] |
| 24 | RIN2 | [GPRIN2, KCNK3, MYOM1] | [[1, 2, 2], [1, 3, 3], [2, 2, 3]] |
| 25 | LMO7 | [TNFRSF19, LIMCH1] | [[1, 2, 3], [1, 2, 3]] |

**Table 15:** Top 25 fuzzy-logic regulatory models identified – II

|  | Model output | Training fit | Test fit |
| --- | --- | --- | --- |
| 1 | TGFBR2 | 0.70 | 0.70 |
| 2 | RIMBP3B | 0.80 | 0.80 |
| 3 | PPP2R1A | 0.72 | 0.71 |
| 4 | UBQLN2 | 0.78 | 0.77 |
| 5 | RIMBP3C | 0.76 | 0.75 |
| 6 | SYNGR3 | 0.76 | 0.77 |
| 7 | POLE | 0.72 | 0.74 |
| 8 | MAOB | 0.72 | 0.74 |
| 9 | PARVG | 0.72 | 0.71 |
| 10 | WNT3A | 0.73 | 0.71 |
| 11 | GALNT12 | 0.72 | 0.74 |
| 12 | SCN2A | 0.71 | 0.74 |
| 13 | DDX60 | 0.71 | 0.68 |
| 14 | SERPINA5 | 0.72 | 0.75 |
| 15 | RIMBP3 | 0.76 | 0.80 |
| 16 | GPR176 | 0.76 | 0.71 |
| 17 | FLCN | 0.71 | 0.65 |
| 18 | MYO1D | 0.75 | 0.69 |
| 19 | KCNQ1 | 0.70 | 0.65 |
| 20 | CPLX2 | 0.74 | 0.68 |
| 21 | TP53 | 0.71 | 0.79 |
| 22 | PIK3CA | 0.70 | 0.61 |
| 23 | ATRNL1 | 0.74 | 0.64 |
| 24 | RIN2 | 0.70 | 0.81 |
| 25 | LMO7 | 0.76 | 0.63 |

**Table 16:** Top 25 fuzzy-logic regulatory models identified - III

| Model output | adj. p-value (BH) |
| --- | --- |
| TGFBR2 | 5.720849e-04 |
| RIMBP3B | 1.907588e-05 |
| PPP2R1A | 9.497439e-05 |
| UBQLN2 | 4.702329e-05 |
| RIMBP3C | 2.522838e-05 |
| SYNGR3 | 1.702524e-04 |
| POLE | 9.527495e-03 |
| MAOB | 1.862614e-04 |
| PARVG | 1.393662e-05 |
| WNT3A | 1.872238e-04 |
| GALNT12 | 2.539112e-03 |
| SCN2A | 2.451987e-04 |
| DDX60 | 1.250976e-04 |
| SERPINA5 | 7.866536e-05 |
| RIMBP3 | 1.961288e-05 |
| GPR176 | 1.117347e-03 |
| FLCN | 5.681593e-03 |
| MYO1D | 2.346973e-05 |
| KCNQ1 | 2.345625e-04 |
| CPLX2 | 1.763330e-05 |
| TP53 | 2.021138e-03 |
| PIK3CA | 7.431795e-04 |
| ATRNL1 | 3.359898e-05 |
| RIN2 | 1.815780e-04 |
| LMO7 | 2.380319e-04 |

#### Models consolidation – Fuzzy logic-based regulatory network

Combining the derived fuzzy logic models into a consolidated network as previously described, we obtained a network with 402 nodes, 849 edges, and a network mean clustering coefficient of 0.018. Consisting predominantly of out-degrees, the maximum degree of 20 is observed at the TP53 gene feature node, followed closely by the SRC (degrees = 19), LONRF2 (degree = 12), PIK3CA (degree = 11), AKT1 (degree = 11), NTN1 (degree = 11), MAPK3 (degree = 10), AURKA (degree = 10), CCND1 (degree = 10), UNC5A (degree = 10), CHEK2 (degree = 10), ICAM1 (degree = 10) and the UBC (degree = 10) gene feature nodes. Average number of neighbors is 3.826, network diameter is 22, characteristic path length is 6.816, and network density is 0.005. Fig. 14

**Figure 14:**
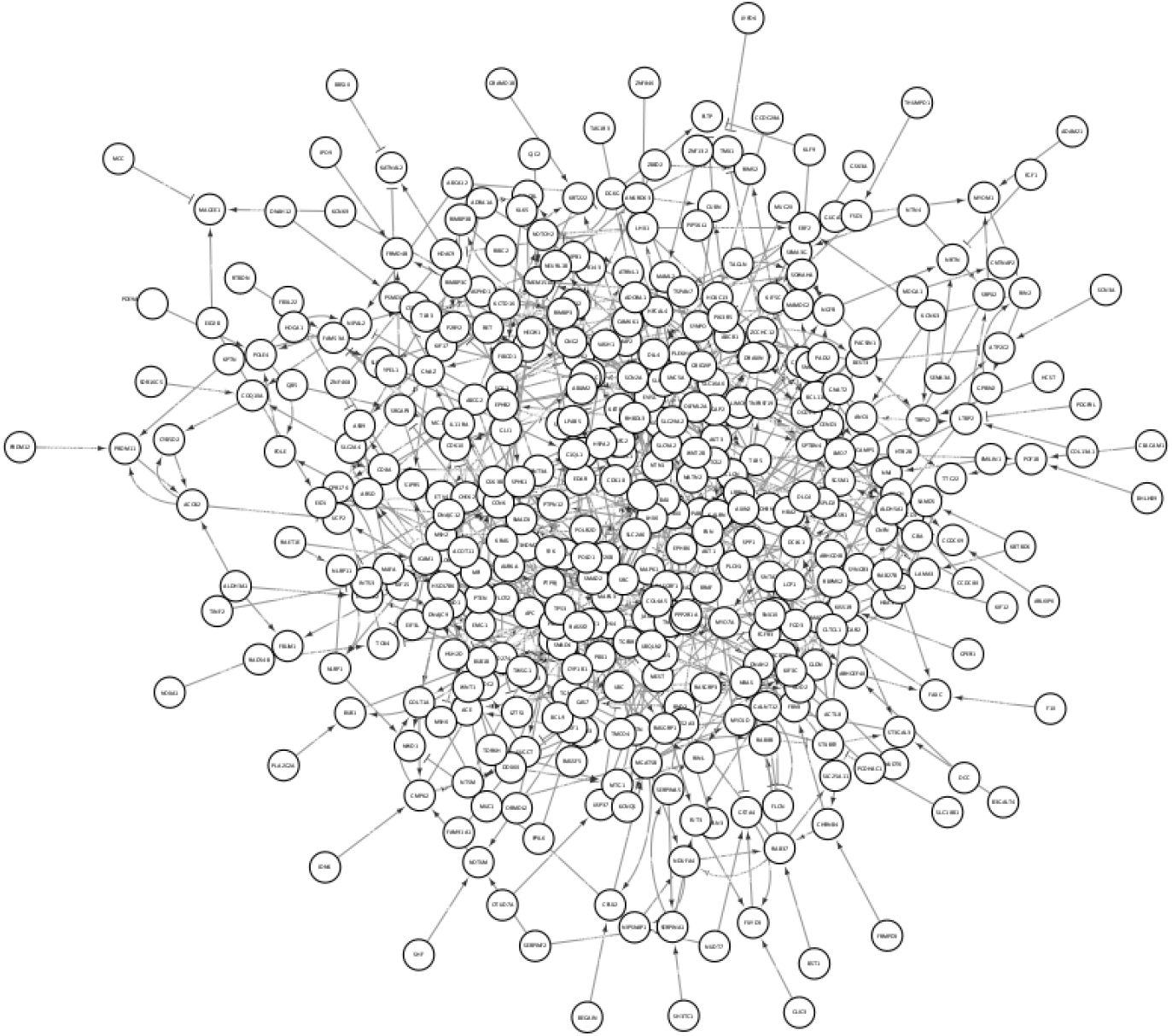
Consolidated fuzzy logic-based regulatory network

### Regulatory Network Validation

Validating regulatory network in independent dataset by topological changes observed on dynamic simulations of inferred regulatory network, no statistical difference is seen in the distribution of monotonic (*t* statistic = –1.5104, p-value = 0.1315) and the adaptive changes (*t* statistic = 1.2079, p-value = 0.2278) across all features over a 5, 000 time steps (Fig. 15 and 16).

**Figure 15:**
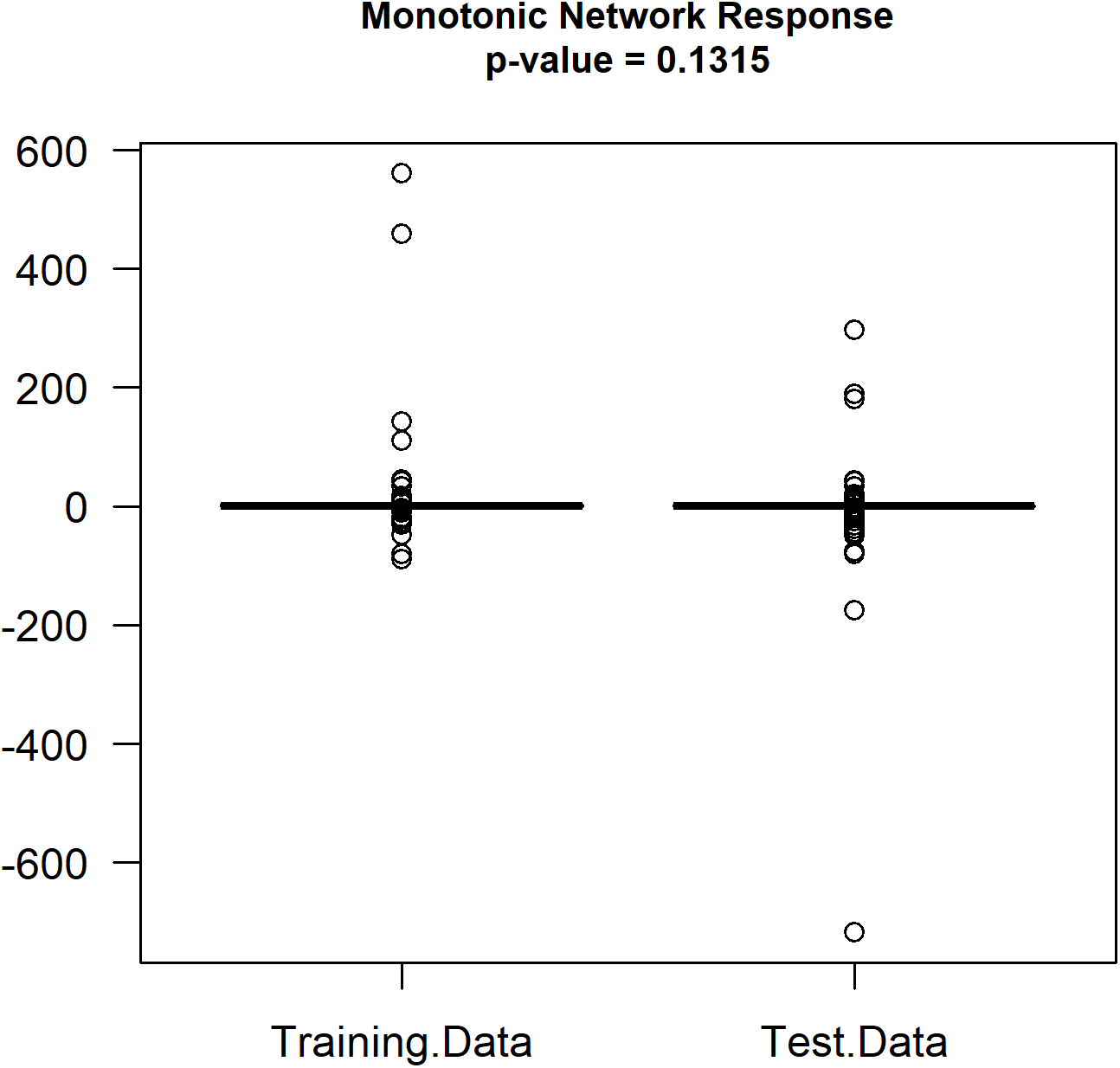
Distribution of observed monotonic changes for all network features over a 5, 000 time step dynamic simulation

**Figure 16:**
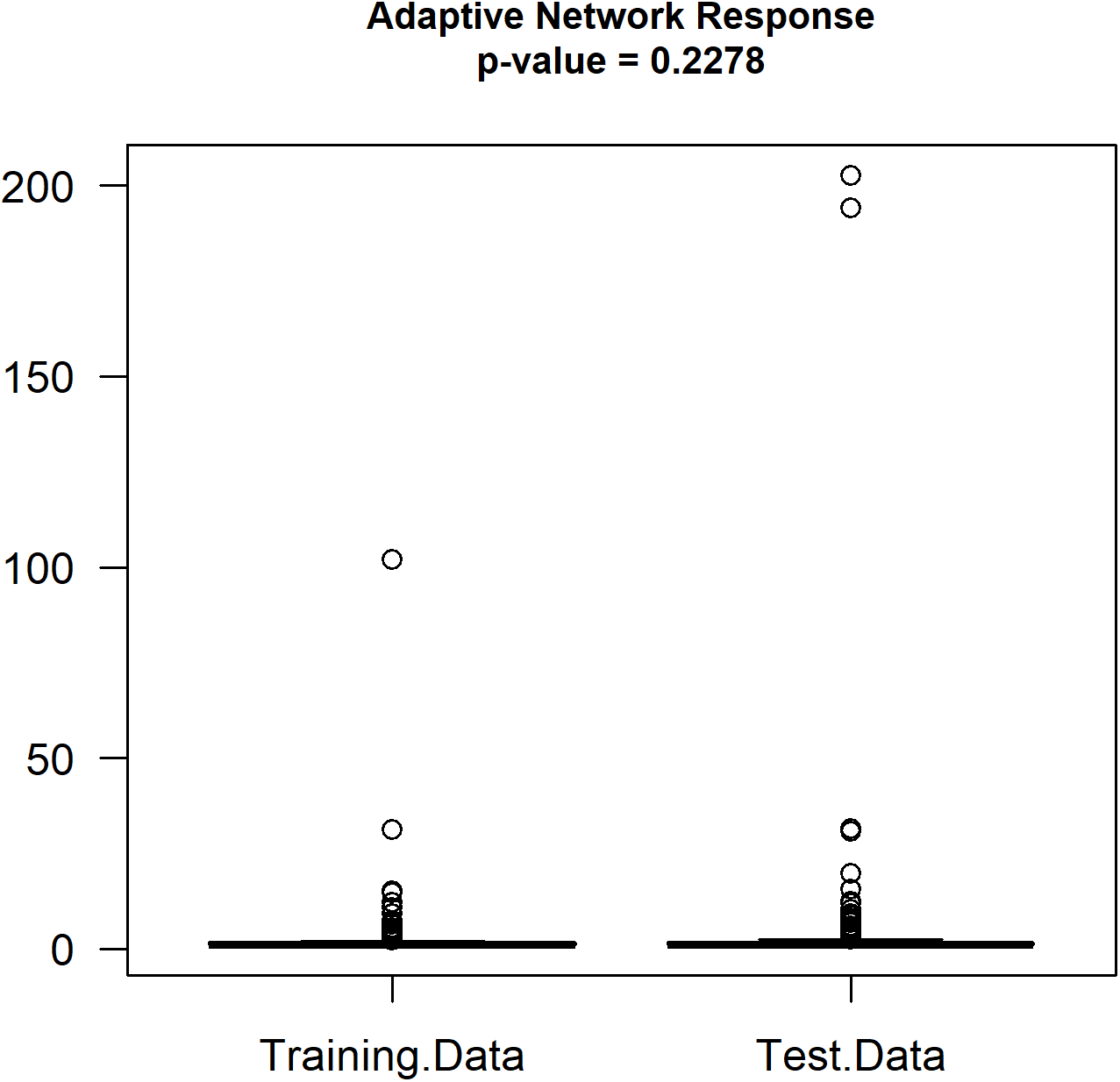
Distribution of observed adaptive changes for all network features over a 5, 000 time step dynamic simulation

### Clinical Significance Evaluation

#### Node importance estimation

Based on the characterized fuzzy logic regulatory network, we computed the hubscore, entropy, mean edge confidence, the associated model fit and the delta change (in fit estimate between training and independent test data), for each gene in the regulatory network (see Methods section) to measure the node importance and to identify key genes driving changes to sensitivity or resistance to Vorinostat in colorectal cancer. Emerging tops with respect to the hubscore, entropy, mean edge confidence, model fit and the delta change scores are the genes TP53, MAGEE1, POLE, TGFBR2 and BUB1 respectively (Supplementary Table). We calculated the normalized score accordingly, to estimate the importance score for each gene (see Methods). Top ranked genes include UBC, PTEN, SMAD2, LMO7, GNAZ, POLR2D, TP53, AKT1, RIMBP3, and CCNK (Table 17).

**Table 17:** Top 40 ranked regulatory features by node importance score estimates.

| Gene Symbol | Description | Rankings |
| --- | --- | --- |
| UBC | ubiquitin C | 1 |
| PTEN | phosphatase and tensin homolog | 2 |
| SMAD2 | SMAD family member 2 | 3 |
| LMO7 | LIM domain 7 | 4 |
| GNAZ | G protein subunit alpha z | 5 |
| POLR2D | RNA polymerase II subunit D | 6 |
| TP53 | tumor protein p53 | 7 |
| AKT1 | AKT serine/threonine kinase 1 | 8 |
| RIMBP3 | RIMS binding protein 3 | 9 |
| CCNK | cyclin K | 10 |
| TNS1 | tensin 1 | 11 |
| PSMD13 | proteasome 26S subunit, non-ATPase 13 | 12 |
| PXN | paxillin | 13 |
| RIMBP3B | RIMS binding protein 3B | 14 |
| RIMBP3C | RIMS binding protein 3C | 15 |
| APC | APC regulator of WNT signaling pathway | 16 |
| GALNT12 | polypeptide N-acetylgalactosaminyltransferase 12 | 17 |
| MAPK1 | mitogen-activated protein kinase 1 | 18 |
| PARVG | parvin gamma | 19 |
| CTNNB1 | catenin beta 1 | 20 |
| MAPK3 | mitogen-activated protein kinase 3 | 21 |
| SMAD3 | SMAD family member 3 | 22 |
| MAGEE1 | MAGE family member E1 | 23 |
| WNT3A | Wnt family member 3A | 24 |
| RASGRP1 | RAS guanyl releasing protein 1 | 25 |
| SERPINA1 | serpin family A member 1 | 26 |
| GPR176 | G protein-coupled receptor 176 | 27 |
| TP53I3 | tumor protein p53 inducible protein 3 | 28 |
| SRC | SRC proto-oncogene, non-receptor tyrosine kinase | 29 |
| DDX60 | DEXD/H-box helicase 60 | 30 |
| POLD1 | DNA polymerase delta 1, catalytic subunit | 31 |
| POLE | DNA polymerase epsilon, catalytic subunit | 32 |
| GSK3B | glycogen synthase kinase 3 beta | 33 |
| TGFBR2 | transforming growth factor beta receptor 2 | 34 |
| KISS1R | KISS1 receptor | 35 |
| HSPG2 | heparan sulfate proteoglycan 2 | 36 |
| BCL9 | BCL9 transcription coactivator | 37 |
| PTPRJ | protein tyrosine phosphatase receptor type J | 38 |
| MGAT5B | alpha-1,6-mannosylglycoprotein 6-beta-N-acetylglucosaminyltransferase B | 39 |
| BUB1B | BUB1 mitotic checkpoint serine/threonine kinase B | 40 |

#### Survival analysis

We reasoned that features driving resistance to vorinostat are very likely drivers of aggressivenes and therefore of poor patient clinical outcomes. We evaluated the potential clinical significance of these vorinostat-resistance associated features in three different ways using colon cancer transcriptomic and clinical data from the cancer genome atlas (TCGA) assayed samples. From the genome data bank retrieved data 334 patient samples had associated clinical information. Of these, 77 samples have had a survival event. The Median time to event is 334.0 days (Mean = 540.7 days), while maximum time to event is 2821.0 days. At different sampling ratios, the 77 samples were randomly divided into a training and a test subdataset and repeating the sample division at each ratio 100 times. Assigning the node importance score of the network feature expression values, Figure 17 shows the AUC estimates’ distribution in the training data and test data weighted logistic regressions. AUC estimates ranged up to 0.99 and 0.93 in the training and test dataset, suggesting a potential optimal subset. Based on data from the TCGA samples, the top 10 percent (by node importance) of features shows a significant association with the survival probability (log-rank p-value = < 0.0001) of colon cancer patients (Fig. 18) – demonstrating the significant clinical relevance of the top identified genes and their roles in cancer progression.

**Figure 17:**
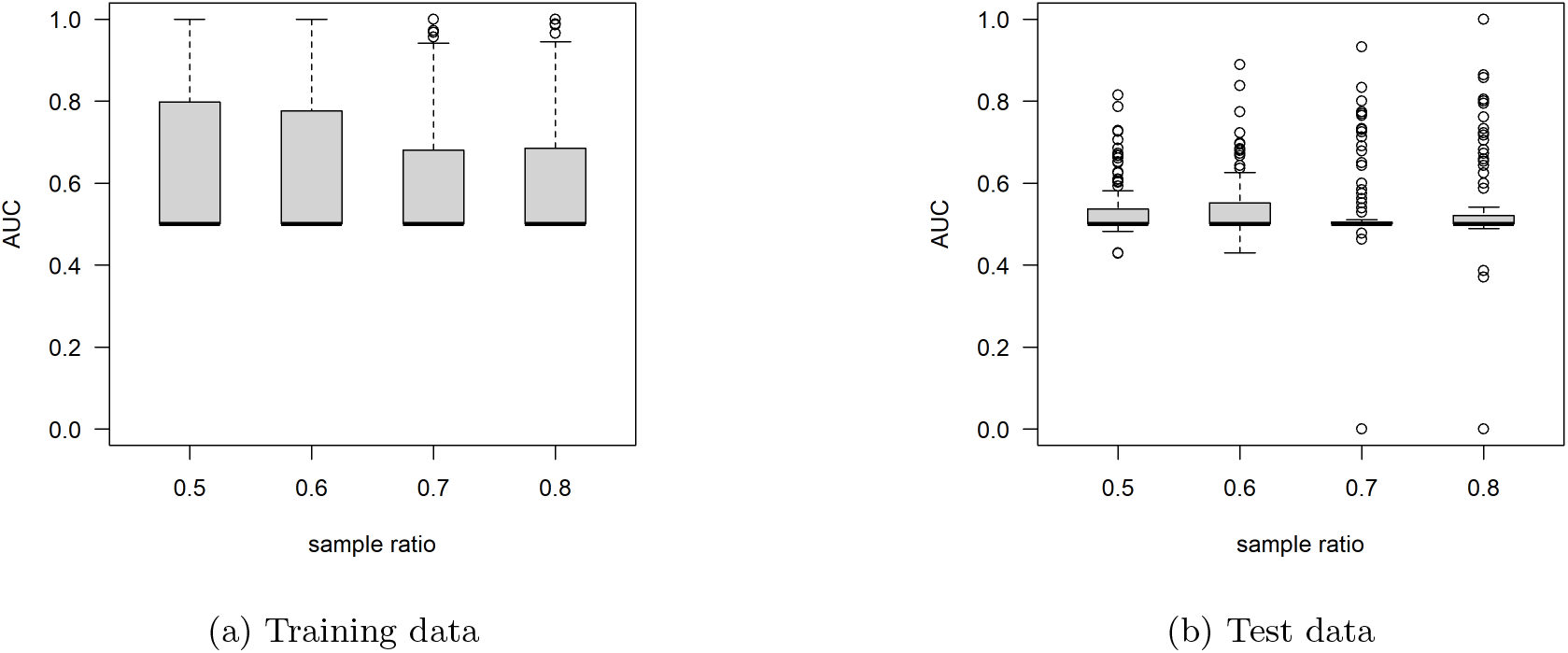
Distribution of AUC estimates in training and test data

**Figure 18:**
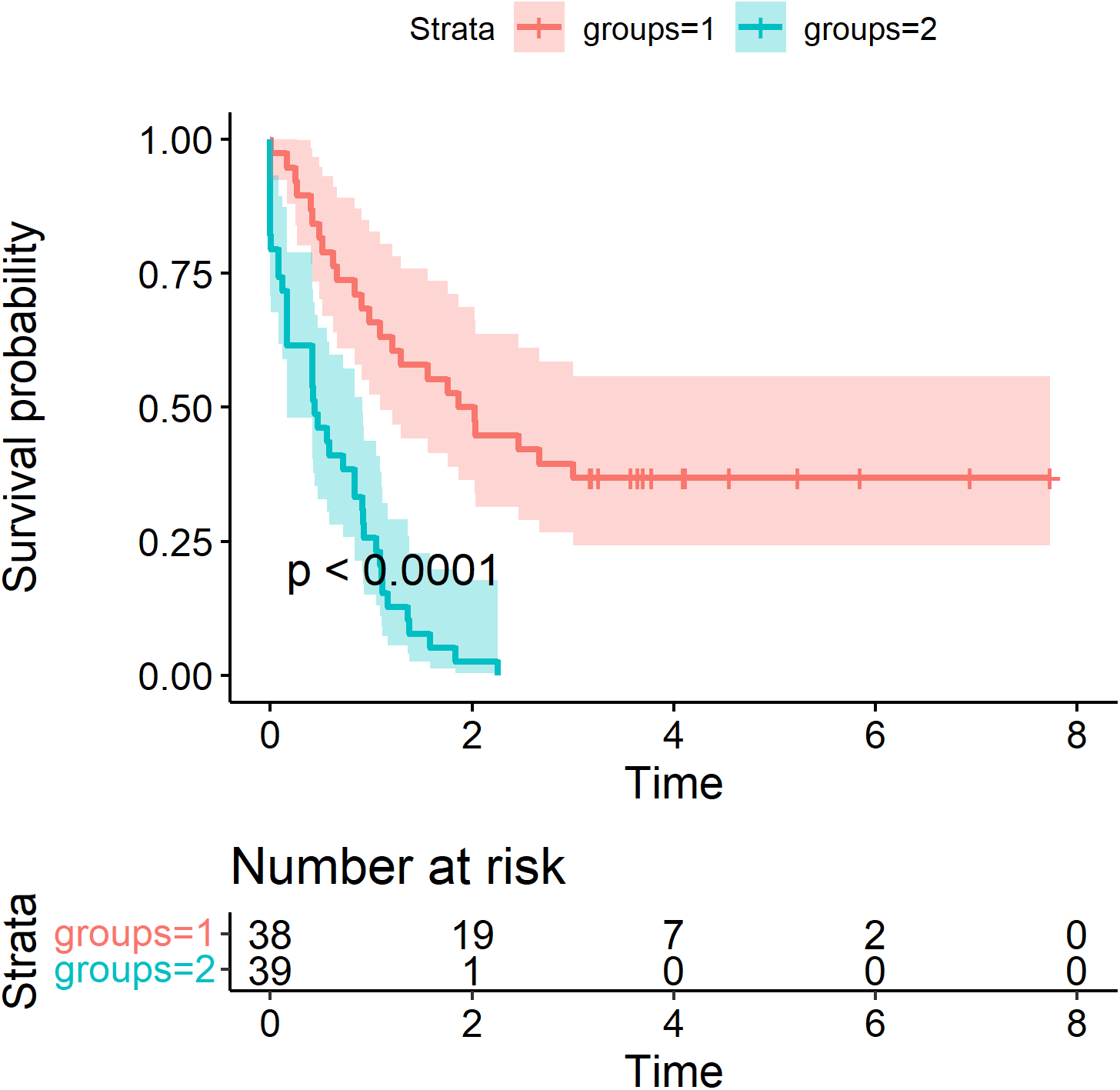
Kaplan-Meier (KM) curve of survival of patients. The groups 1 and 2 were identified by optimal separation of predicted responses from a Cox proportional hazard fit of the survival model

#### Tools and implementations

The fuzzy logic regulatory inference method of Gormley et al [95] and its optimization for quicker time to inference is implemented in the platform independent Java programing language. Its source codes and precompiled binaries are freely available at the repository locations:

- https://github.com/paiyetan/jfuzzymachine
- https://github.com/paiyetan/jfuzzymachine/releases/tag/v1.7.21
- https://bitbucket.org/paiyetan/jfuzzymachine/src/master/
- https://bitbucket.org/paiyetan/jfuzzymachine/downloads/

All statistical analysis was done in the R programing and computational statistics enviromment [59]. Network diagrams and analysis were also done using the Cytoscape tool, version 3.8.2 running on a Mac OS X 10.15.7 - x86_64 operating system.

## Discussions

Though the prevalence of colon cancer appear to be on the decline particularly in the above 65 year olds, the rising incidence in the younger population remain of significant concern. Besides surgical excision of tumor tissue and the use of classical chemotherapeutic regiments such as 5-FU, oxaliplatin, irinotecan, capecitabine, leucovorin, etc. with or without radiation, the search for more rational therapy continue to gain traction. The approval of bevacizumab, panitumumab, and cetuximab for colorectal cancer buttress the potential clinical benefit of rationally designed therapies – targeted therapies developed to take advantage of unique alterations specific to cancer cells to maximize the desired therapeutic effect in malignant cells and minimize the toxicity in normal cells [121]. Currently approved for metastatic, stage IV or recurrent disease, other targeted therapies for colorectal cancer include aflibercept, ramucirumab, panitumumab, regorafenib, and pembrolizumab. These are designed against the VEGF, EGFR, or tyrosine kinases. Targeting the VEGF pathway, bevacizumab and ramucirumab are developed as monoclonal antibodies while aflibercept, a recombinant fusion protein. Cetuximab and panitumumab, targets the EGFR pathway upstream of KRAS, and particularly effective in non-KRAS activating mutation cancers. More recently approved based on on data from 149 patients with MSI-H or DNA mismatch repair cancers enrolled across 5 uncontrolled, multicohort, multicenter, single-arm clinical trials, Pambrolizumab is an antibody that targets the programmed cell death 1 (PD-1) protein in patients with microsatellite unstable tumors – associated with germline defects in the MLH1, MSH2, MSH6, and PMS2 genes[122, 123]. The proposed relationship of pambrolizumab to the DNA mismatch repair genes or gene products (MLH1, MSH2, MSH6, and PMS2) is less direct, compared to that of bevacizumab, ramucirumab or aflibercept to VEGF. Pambrolizumab targets the PD-1 protein on T cells, preventing its association with the PD-L1 ligand on tumor cells, macrophages or other tumor-infiltrating lymphocytes and myeloid cells acting in concert to suppress T-cells activation[124]. T cell activation results from presentation of mutation-associated neoantigen (MANA) that results from protein products consequent to DNA mismatches in complex with the major histocompatibility complex (MHC) protein [125].

Analogous to pambrolizumab and other targeted therapies, vorinostat can act in both direct and indirect ways on the molecular pathways regulating oncogenic processes. In addition to inhibiting histone deacetylases, evidences also point to its effect on the posttranslational modification state of proteins involved in oncogenic and tumor supression processes. Vorinostat inhibits the removal of acetyl group from the *ϵ*-amino group of lysine residues of histone proteins by histone deacetylases (HDACs) maintaining chromatin in an expanded state and thus facilitating transcriptional activities of major regulatory gene products such as transcription factors, cell-signaling regulatory proteins, and proteins regulating cell death[35]. Particularly described is its effect on the cyclin-dependent kinase inhibitor 1A, CDKN1A (also known as p21, WAF1/CIP1) gene transcription[36]. Richon et al found that vorinostat selectively induces CDKN1A expresion[36]. By binding to cyclin dependent kinases, CDKN1A prevents the phosphorylation of cyclin-dependent kinase substrates and thus block cell cycle progression[126].

Non-histone-protein related effect of vorinostat include increased DNA binding of the transcriptional activator TP53 (Tumor protein 53, also known as p53) from increased acetylation, and a consequent increase in p53-regulated gene transcription rate. Also, BCL6 repression of transcription is inhibited by increased acetylation as a result of vorinostat[127]. As opposed to increased transcription, vorinostat (HDACi) represses the expression of genes cyclin D1, ErbB2, and thymidylate synthase [128]. Evidently, the effect of vorinostat tips the cellular equilibriun toward cell cycle arrest, anti-proliferation and apoptosis. It thus appeal to reason that molecular pathways of resistance would be quite the opposite. In attempts to circumvent resistance, Falkenberg et al had through a functional genomics screen identified genes that when knocked down by RNA interference (RNAi) sensitized cells to vorinostat-induced apoptosis. In other words, when these genes are knocked down, they co-operated with vorinostat to induce tumour cell apoptosis in otherwise resistant cells (synthetic lethality). These included – BEGAIN, CCNK, CDK10, DPPA5, EIF3L, GLI1, JAK2, NFYA, POLR2D, PSMD13, RGS18, SAP130, TGM5, and TOX4 (see Table 18)[39], all of which are pro-proliferative and potentially oncogenic. Of importance to our study however is to determine the molecular processes underneath the observed synthetic lethal phenotype and potential clinical significance. In a follow-up study, Falkenberg et al had validated the GLI1 gene as co-operative with vorinostat[40] to induce cell cycle arrest and apoptosis in otherwise vorinostat-resistant colon cancer cell lines. Given GLI1’s role in the sonic hedgehog pathway, we had hypothesized that resistance to vorinostat is a result of uptick in embryonal gene regulatory programs. We also hypothesized that elucidated regulatory mechanism would include crosstalks that regulate this biological processes – embryonal gene regulatory programs.

**Table 18:**
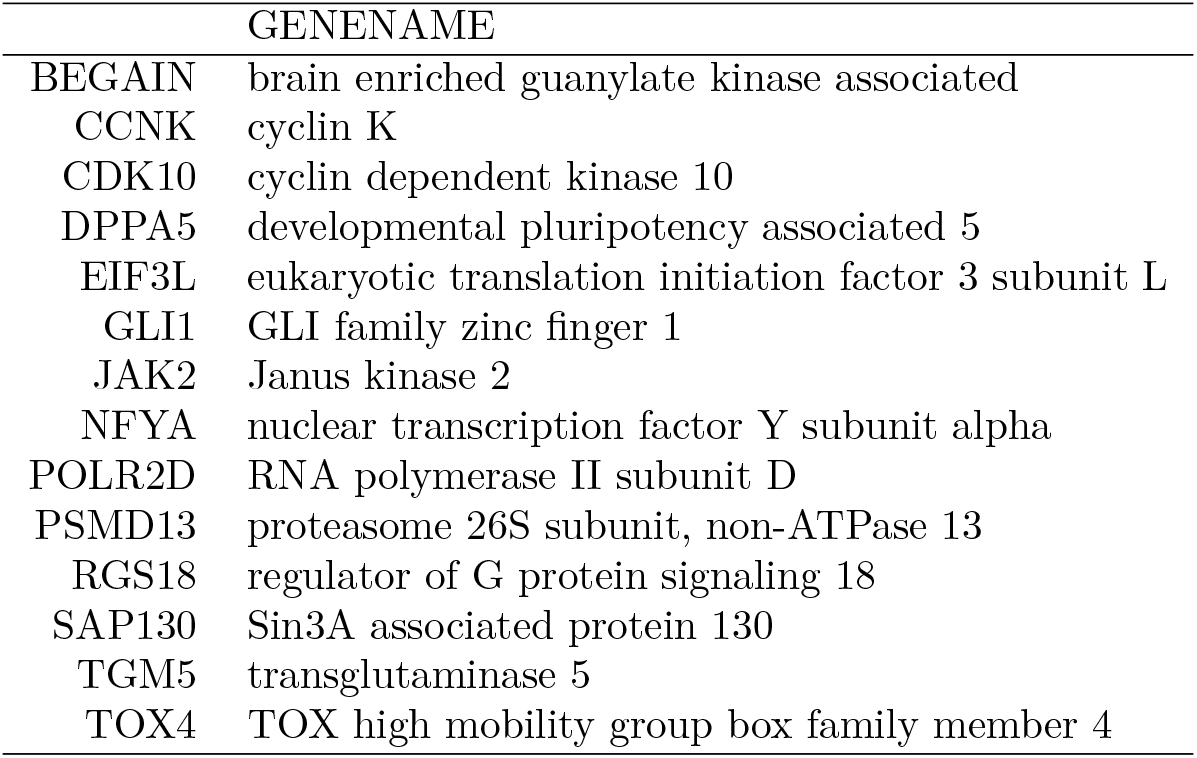
Table of identified synthetical lethal gene partners to histone deacetylase by Falkenberg et al.

Glioma-Associated Oncogene Homolog 1 (GLI1) is a zinc finger protein and a transcription factor that acts downstream of the Hedgehog (Hh) signaling pathway. It mediates morphogenesis, cell proliferation and differentiation[129–132]. From reports of its potential relationship to GLI1 and vorinostat, Falkenberg and collagues reported the repression of the BCL2L1 gene on GLI1 knockdown. Bcl-2-Like Protein 1 (BCL2L1) is a cell death inhibitor, it inhibits the activation of caspases by binding to and blocking the voltage-dependent anion channel (VDAC), preventing the release of the caspase activator, CYC1, from the mitochondrial membrane[134, 135]. At a adjusted p-value of < 0.05, we found the differential expression of BCL2L1 to significantly change in up to 8 siRNA knockdown (synthetic lethality) experiments, including siGLI1 knockdown (Adjusted p-value = 1.7444e–21, Log2 fold change = –0.6694, St. error = 0.0676), compared to controlled experiments. However, subject to our selection criteria, BCL2L1 did not make the list of selected features for regulatory network inference. Although it may not be generalized in this study, there is evidence of the potential utility of BCL2L1 repression consequent to GLI1 knockdown as a path to restoring sensitivity to vorinostat in resistant colon cancer cell lines.

The Sonic Hedgehog (SHH) pathway upstream of GLI1 consists of PTCH1, SMO, and SUFU. Binding of the Hedgehog ligand to the cell surface receptor Patched (PTCH) releases its inhibitory effect on Smoothened (SMO), which in turn activates GLI1 [40, 136]. Suppressor Of Fused Homolog, SUFU down-regulates transactivation of target genes by GLI1 [137, 138]. It forms a part of the co-repressor complex that acts on DNA-bound GLI1 and may act by linking GLI1 to BTRC – targeting GLI1 for degradation by proteasome [130, 138–140]. Amongst TTRUSTv2 transcription factor-target database [141] retrieved 19 gene-targets of GLI1, only 2, AKT1[142] and SMAD4 [143] are contained in the derived fuzzy-logic regulatory network. Both of these are also known to be regulated by PIK3A and TGF-*β* respectively. Argawal et al. and, Nye et al. had suggested a form of cross-talk between the Sonic Hedgehog pathway and the cell proliferative pathways of AKT1 and TGF-*β*, mediated through GLI1 and GLI1-SMAD4 complex respectively. However, the non-significance of other members of the SHH pathway in our derived fuzzy logic regulatory network questions its role in vorinostat-resistance or re-sensitivity in the HCT116 colorectal cancer cell lines. The significance of other canonical upstream regulators of AKT1 and SMAD4, PIK3CA (node importance rank = 77; model fit =0.701, model adjusted p-value = 0.00074) and TGFBR2 (node importance rank = 34, model fit = 0.704, model adjusted p-value = 0.00057) respectively suggests alternate mechanisms of resistance or re-sensitivity (Figs 19 and 20). These interactions may in part explain the proliferative and anti-proliferative processes observed in vorinostat resistant and re-sensitized colon cancer cell lines independent of the SHH pathway. In some form multiple feed-forward (FF) and positive feed-back (PFB) control manner, GLI1 on the other hand appears to be under regulatory control with RET and ETV4, which themselves are tightly regulated by NEURL1B and AURKA. A consequently amplified GLI1 signal activates ABLIM2, whose signal is tempered by SRC. (Fig. 21). Using in-vivo cell culture and xenograft models, Ruan et al. recently showed that RET (rearranged during transfection) enhanced transcriptional activation by HH, independent of the SHH pathway. They showed that inhibition of GLI1 led to a reduction of RET-induced proliferation of SH-SY5Y cells and outgrowth of xenografts[144]. The role of GLI1 on RET expression in neuroblastoma is well documented – GLI1 induces the expression of RET[145, 146]. Zhu et al in a xenograft model, showed that EVS variant transcription factor 4 (ETV4) depletion inhibits the CXCR4/SHH/GLI1 signaling cascade in breast cancer[147]. Such may yet be the case in the vorinostat resistant colorectal cancer cell lines.

**Figure 19:**
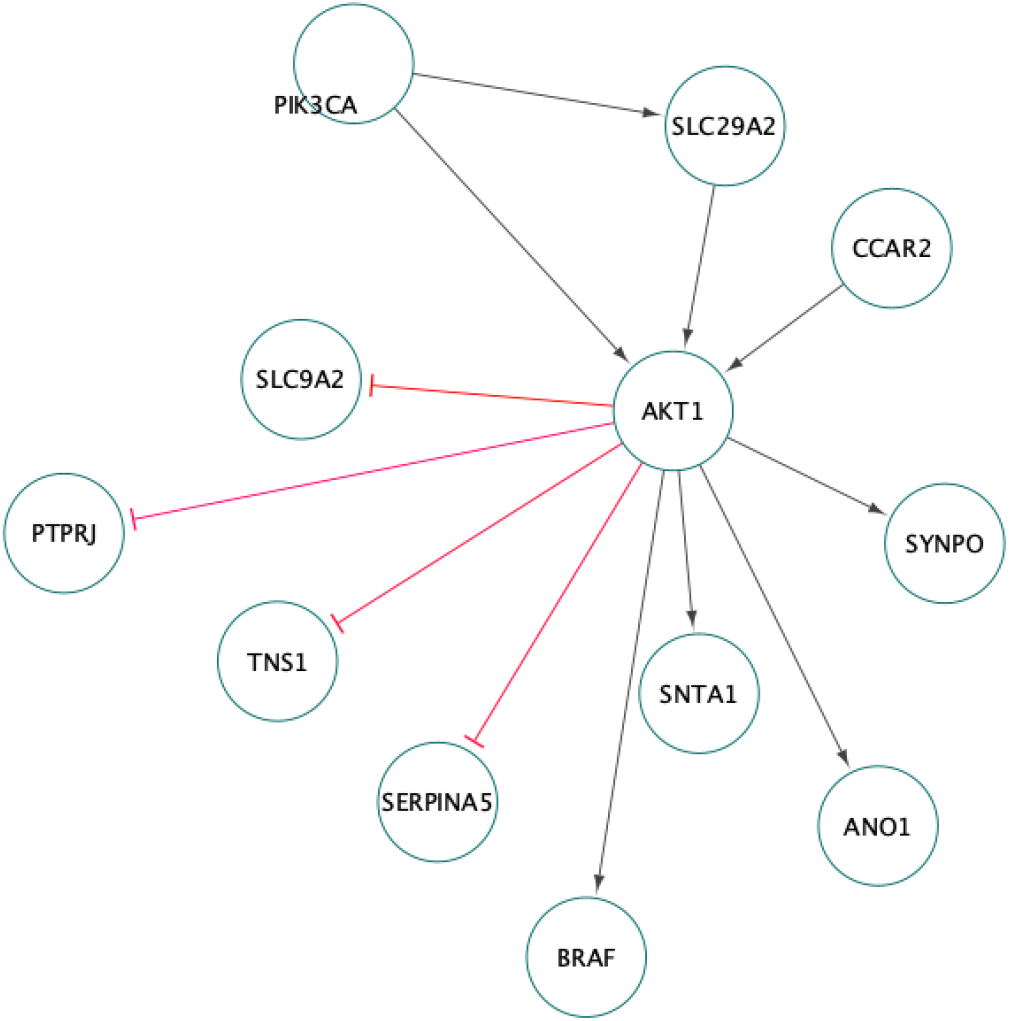
The AKT1 Pathway. The PIK3CA-AKT1-BRAF relationship remains consistent as with canonical cell pro-survival and proliferation pathway. Besides less known activation pathways involving dowstream activation of SNTA1, ANO1 and SYNPO, the canonical AKT1 activation of BRAF is highlighted by the fuzzy logic inference method. PIK3CA upregulation of ANO1 through AKT1 may be a mechanism of resistance to circumvent vorinotat’s activity, particularly in response to GLI1-knockdown (see text).

**Figure 20:**
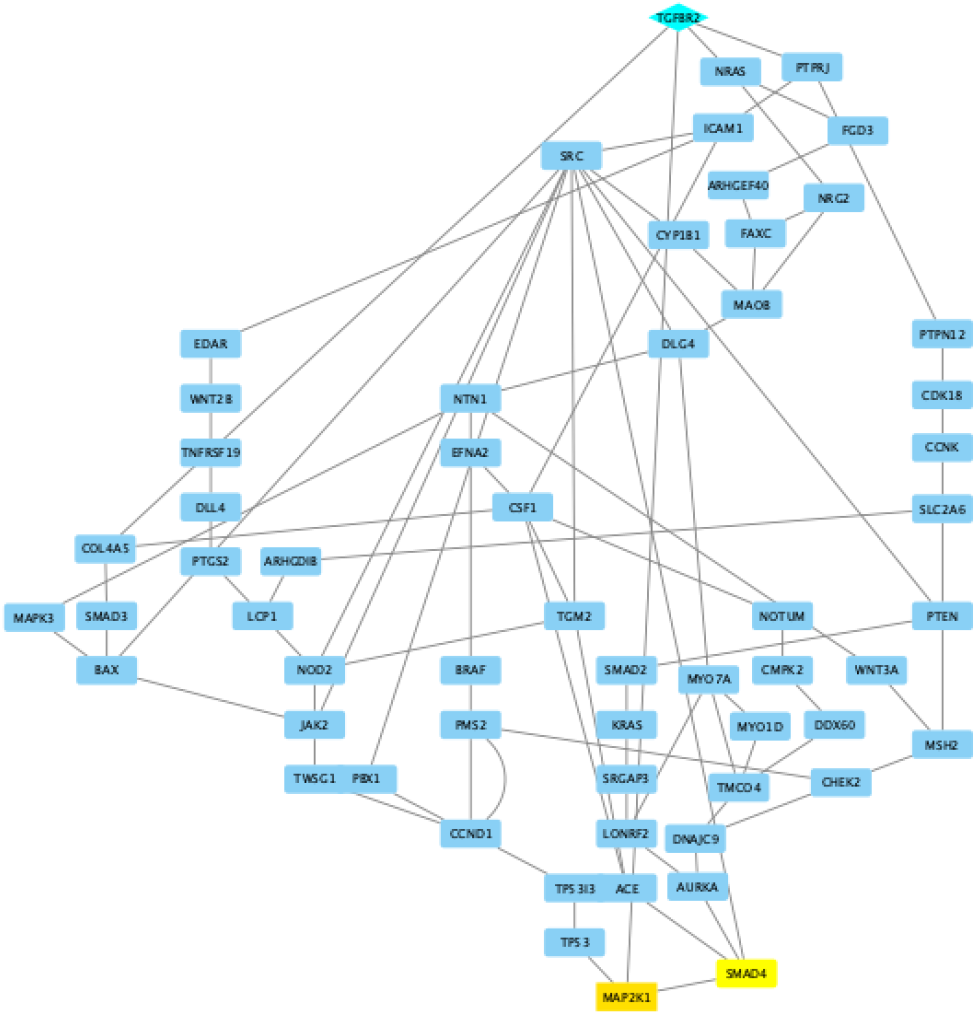
The TGFBR2-SMAD4 subnetwork. The Fuzzy logic based network inference approach shows canonical and non-canonical interactions that connect the TGFBR2 to SMAD4. Canonically, activated carboxy-terminal phosphorylated SMADs (SMAD2 and SMAD3) partner with their common signal transducer SMAD4 and translocate into the nucleus to regulate diverse biological activites, mostly by partnering with transcription factors. Inferred fuzzy-logic regulatory network includes both direct and indirect relationships amongst gene and gene-products related to the SMAD signaling complex.

**Figure 21:**
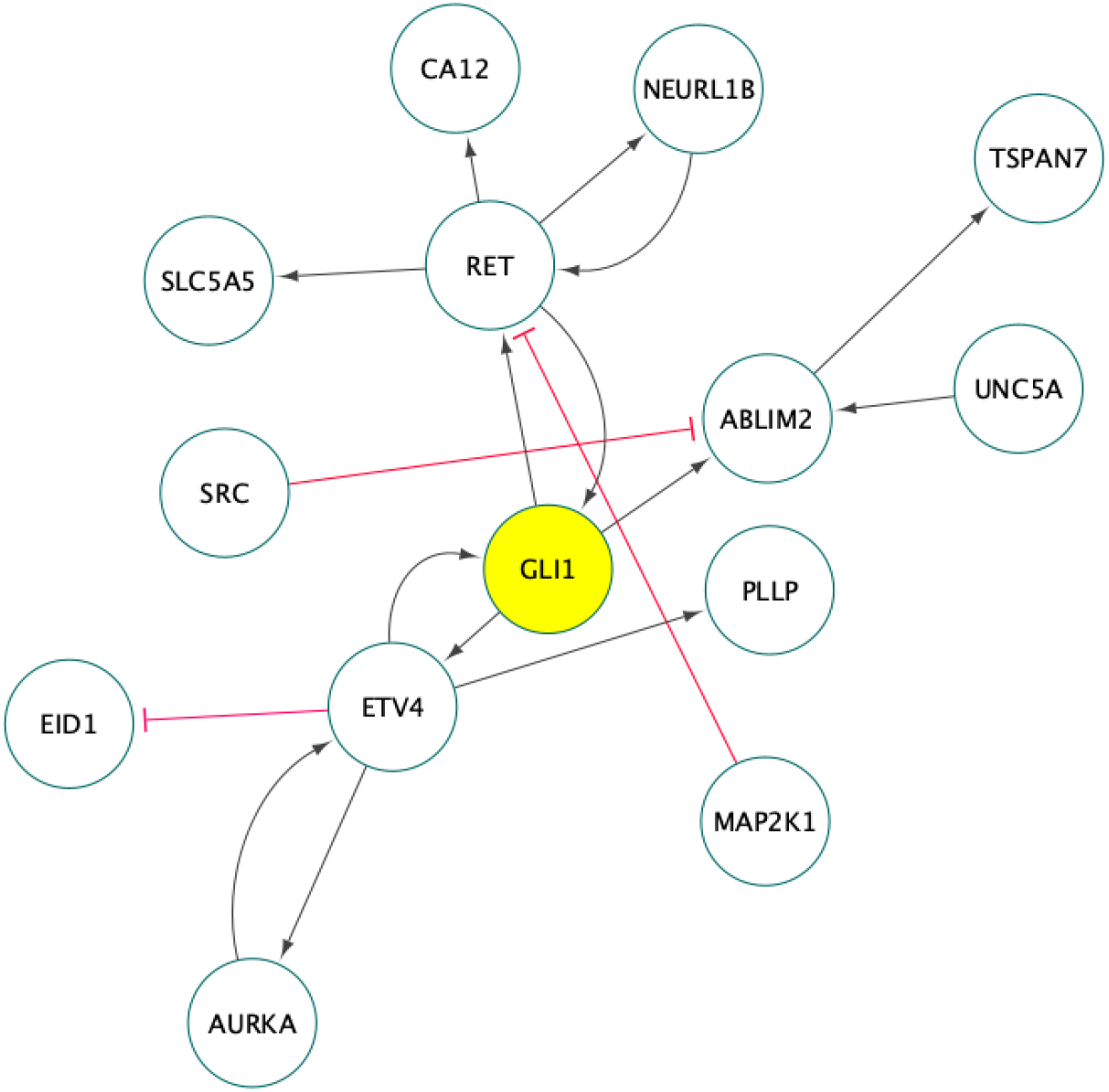
Inferred GLI1 interactions based on fuzzy logic. GLI1 activation or regualation appears to be independent of members of the Sonic Hedgehog (SHH) pathway. In some form multiple feed-forward (FF) and positive feed-back (PFB) control manner, GLI1 appears to be under regulatory control with RET and ETV4, which themselves are tightly regulated by NEURL1B and AURKA respectively. Amplified GLI1 signal activates ABLIM2, whose signal is tempered by SRC.

PIK3CA upregulation of anoctamin 1 (ANO1) through AKT1 may be a mechanism of resistance to circumvent vorinotat’s activity, particularly in response to GLI1 knockdown. Mazzone et al [148], using a luciferase reporter system to determine ANO1 promoter activity, chromatin immunoprecipitation, siRNA knockdown, PCR, immunolabeling, and recordings of Ca^2+^-activated Cl^-^currents in human embryonic kidney 293 (HEK293) cells showed that binding of GLI1 represses ANO1 expression. They also showed that knocking down of GLI1 expression and inhibition of its activity increased the expression of ANO1 transcripts and Ca^2+^-activated Cl^-^currents in HEK293 cells. Relating to the activity of PIK3CA, Mroz and colleagues [149] showed that induction of the transmembrane protein 16A (TMEM16A) also known as ANO1 expression is mediated by a sequential activation of phosphatidylinositol 3-kinase (PIK3) and protein kinase C-*δ* (PKC*δ*). Our fuzzy logic approach indicates an involvement of AKT (Fig. 19). We suppose that in response to vorinostat, the PIK3CA-AKT-ANO1 pathway provides alternate escape pathway from anti-cell profliferation signatures.

With evidences pointing towards activation of cell pro-survival and cell proliferation pathways independent of SHH, the role of these pro-survival and cell proliferation pathway members (PIK3CA, AKT1, MAPK1, MAPK3, WNT3A etc) in vorinostat resistance, restoring vorinostat sensitivity or improving patient clinical response in characteristic colorectal cancer, are worth evaluating. Interestingly observed are the almost equal or more significant represention of tumor suppressor genes (PTEN, TP53, APC, UBC, GSK3B, etc) top-ranked in terms of node importance in the derived regulatory network – very likely playing the role to restore vorinostat sensitivity in the vorinostat-resistant colorectal cancer cell lines. These relationships appear to be clinically significant, given top-ranked features’ expression being predictive of colorectal cancer patients survival (Fig. 18, log rank, p-value < 0.0001) in the sampled population. Here is presented a rationale for including anti-cell prosurvival and anti-cell proliferative genes’ targeted therapy, in combination with vorinostat therapy to improve patient survival in colorectal cancer.

**Figure 22:**
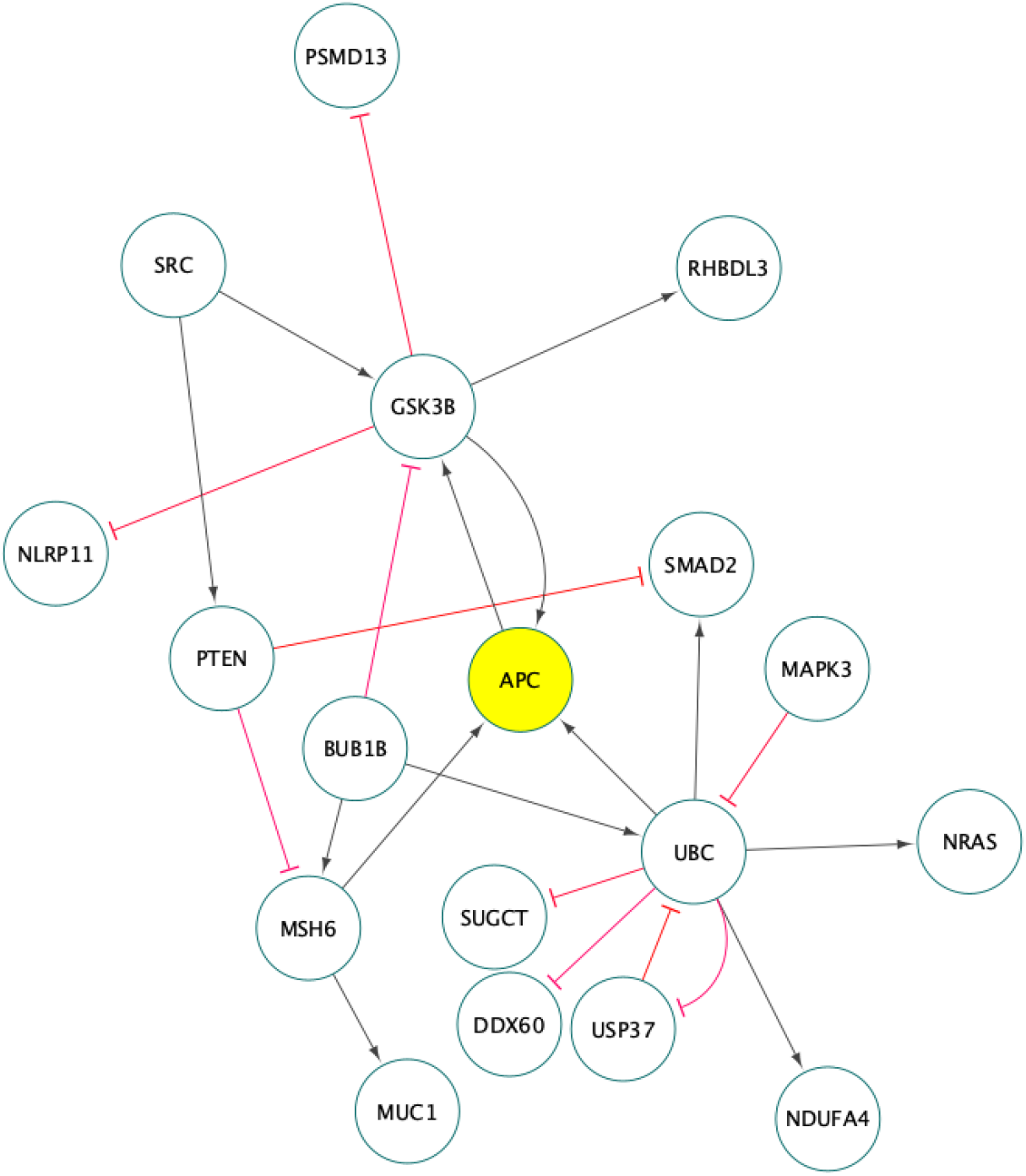
The APC regulation. Remaining consistent is the UBC-APC-GSK3B interaction

## Conclusions

Our knowledge-guided fuzzy logic approach is able to tease the regulatory mechanism involved in histone deacetylase inhibition resistance in colon cancer cell lines from biological dataset.

There is no significant evidence that vorinostat resistance is due to an upregulation of emmbryonal gene regulatory pathways. Our observation rather support a topological rewiring of canonical oncogenic (pro-cell survival, cell proliferative) pathways, including the PIK3CA, AKT, RAS, BRAF etc. pathways. Exploring the potential clinical or biomedical significance, inferred major regulatory molecules are able to delineate patients into high- and low-risk of mortality. The identified key regulatory network genes’ expression profile are able to predict short-to medium-term survival in colorectal cancer patients – providing a rationale for an effective combination of therapeutics that target these genes (particularly the pro-cell survival and cell proliferative gene products) along with vorinostat in the treatment of colorectal cancer.

## Study Limitations

Apparent is the paucity of siRNA knockdown data to encompass all potentially possible epistatic interactions that may be associated with re-sensitizing vorinostat-resistant colorectal cancer. The fuzzy logic approach models direct and indirect interactions which inherently associates relationship in molecular abundance with regulatory activation or inhibition; it incorporates no epigenetic, trans- or cis-genomic signatures independent of abundance and the consequent effects in the regulatory models inferred.

## Acknowledgements

A special thanks to Dr. Iosif Vaisman, Dr. Dmitri Klimov, and Dr. Saleet Jafri for their advises on the greater body of work presented here. Here is also acknowledging the high performance computing resource at the Frederick National Laboratory for Cancer Research (FNLCR) and the Extreme Science and Engineering Discovery Environment (XSEDE) supported by the National Science Foundation at the Texas Advanced Computing Center (TACC) at The University of Texas at Austin - http://www.tacc.utexas.edu. The Frederick National Laboratory for Cancer Research is wholly funded by the National Cancer Institute.

## References

1. Alliance CC. Colorectal cancer, know the facts. 2021. https://www.ccalliance.org/colorectal-cancer-information/facts-and-statistics.

2. Society AC. Key statistics for colorectal cancer. 2021. https://www.cancer.org/cancer/colon-rectal-cancer/about/key-statistics.html.

3. Siegel RL, Miller KD, Goding Sauer A, Fedewa SA, Butterly LF, Anderson JC, et al. Colorectal cancer statistics, 2020. CA: a cancer journal for clinicians. 2020;70:145–64.

4. Disease Control C for, Prevention. United states cancer statistics: Data visualizations, leading cancer (colon and rectum) cases and deaths, all races/ethnicities, male and female, 2017. 2021. https://gis.cdc.gov/Cancer/USCS/DataViz.html.

5. Society AC. Colorectal cancer risk factors. 2021. https://www.cancer.org/cancer/colon-rectal-cancer/causes-risks-prevention/risk-factors.html.

6. Mármol I, Sánchez-de-Diego C, Pradilla Dieste A, Cerrada E, Rodriguez Yoldi MJ. Colorectal carcinoma: A general overview and future perspectives in colorectal cancer. International journal of molecular sciences. 2017;18:197.

7. Fakih MG, Pendyala L, Fetterly G, Toth K, Zwiebel JA, Espinoza-Delgado I, et al. A phase i, pharmacokinetic and pharmacodynamic study on vorinostat in combination with 5-fluorouracil, leucovorin, and oxaliplatin in patients with refractory colorectal cancer. Clinical Cancer Research. 2009;15:3189–95.

8. Fakih MG. A phase i, pharmacokinetic, and pharmacodynamic study of two schedules of vorinostat in combination with 5-fluorouracil and leucovorin in patients with refractory solid tumors. 2010.

9. Ree AH, Dueland S, Folkvord S, Hole KH, Seierstad T, Johansen M, et al. Vorinostat, a histone deacetylase inhibitor, combined with pelvic palliative radiotherapy for gastrointestinal carcinoma: The pelvic radiation and vorinostat (pravo) phase 1 study. The lancet oncology. 2010;11:459–64.

10. Morelli MP, Tentler JJ, Kulikowski GN, Tan A-C, Bradshaw-Pierce EL, Pitts TM, et al. Preclinical activity of the rational combination of selumetinib (azd6244) in combination with vorinostat in kras-mutant colorectal cancer models. Clinical Cancer Research. 2012;18:1051–62.

11. Fakih M, Groman A, McMahon J, Wilding G, Muindi J. A randomized phase ii study of two doses of vorinostat in combination with 5-fu/lv in patients with refractory colorectal cancer. Cancer chemotherapy and pharmacology. 2012;69:743–51.

12. Vansteenkiste J, Van Cutsem E, Dumez H, Chen C, Ricker JL, Randolph SS, et al. Early phase ii trial of oral vorinostat in relapsed or refractory breast, colorectal, or non-small cell lung cancer. Investigational new drugs. 2008;26:483–8.

13. Wilson PM, El-Khoueiry A, Iqbal S, Fazzone W, LaBonte MJ, Groshen S, et al. A phase i/ii trial of vorinostat in combination with 5-fluorouracil in patients with metastatic colorectal cancer who previously failed 5-fu-based chemotherapy. Cancer chemotherapy and pharmacology. 2010;65:979–88.

14. Deming DA, Ninan J, Bailey HH, Kolesar JM, Eickhoff J, Reid JM, et al. A phase i study of intermittently dosed vorinostat in combination with bortezomib in patients with advanced solid tumors. Investigational new drugs. 2014;32:323–9.

15. Fu S, Hou M, Naing A, Janku F, Hess K, Zinner R, et al. Phase i study of pazopanib and vorinostat: A therapeutic approach for inhibiting mutant p53-mediated angiogenesis and facilitating mutant p53 degradation. Annals of oncology. 2015;26:1012–8.

16. Mahalingam D, Mita M, Sarantopoulos J, Wood L, Amaravadi RK, Davis LE, et al. Combined autophagy and hdac inhibition: A phase i safety, tolerability, pharmacokinetic, and pharmacodynamic analysis of hydroxychloroquine in combination with the hdac inhibitor vorinostat in patients with advanced solid tumors. Autophagy. 2014;10:1403–14.

17. Wang Y, Janku F, Piha-Paul S, Hess K, Broaddus R, Liu L, et al. Phase i studies of vorinostat with ixazomib or pazopanib imply a role of antiangiogenesis-based therapy for tp53 mutant malignancies. Scientific reports. 2020;10:1–9.

18. Grant S, Easley C, Kirkpatrick P. Vorinostat. Nature reviews Drug discovery. 2007;6:21–2.

19. Stowell JC, Huot RI, Van Voast L. The synthesis of n-hydroxy-n’-phenyloctanediamide and its inhibitory effect on proliferation of axc rat prostate cancer cells. Journal of medicinal chemistry. 1995;38:1411–3.

20. Richon V, Webb Y, Merger R, Sheppard T, Jursic B, Ngo L, et al. Second generation hybrid polar compounds are potent inducers of transformed cell differentiation. Proceedings of the National Academy of Sciences. 1996;93:5705–8.

21. Butler LM, Agus DB, Scher HI, Higgins B, Rose A, Cordon-Cardo C, et al. Suberoylanilide hydroxamic acid, an inhibitor of histone deacetylase, suppresses the growth of prostate cancer cells in vitro and in vivo. Cancer research. 2000;60:5165–70.

22. Huang L, Pardee AB. Suberoylanilide hydroxamic acid as a potential therapeutic agent for human breast cancer treatment. Molecular medicine. 2000;6:849–66.

23. Munster PN, Troso-Sandoval T, Rosen N, Rifkind R, Marks PA, Richon VM. The histone deacetylase inhibitor suberoylanilide hydroxamic acid induces differentiation of human breast cancer cells. Cancer research. 2001;61:8492–7.

24. Cooper AL, Greenberg VL, Lancaster PS, Nagell Jr JR van, Zimmer SG, Modesitt SC. In vitro and in vivo histone deacetylase inhibitor therapy with suberoylanilide hydroxamic acid (saha) and paclitaxel in ovarian cancer. Gynecologic oncology. 2007;104:596–601.

25. Krug LM, Curley T, Schwartz L, Richardson S, Marks P, Chiao J, et al. Potential role of histone deacetylase inhibitors in mesothelioma: Clinical experience with suberoylanilide hydroxamic acid. Clinical lung cancer. 2006;7:257–61.

26. Miles Prince H, Bishton M, Harrison S. The potential of histone deacetylase inhibitors for the treatment of multiple myeloma. Leukemia & lymphoma. 2008;49:385–7.

27. Richon VM, Garcia-Vargas J, Hardwick JS. Development of vorinostat: Current applications and future perspectives for cancer therapy. Cancer letters. 2009;280:201–10.

28. Claerhout S, Lim JY, Choi W, Park Y-Y, Kim K, Kim S-B, et al. Gene expression signature analysis identifies vorinostat as a candidate therapy for gastric cancer. PloS one. 2011;6:e24662.

29. Galanis E, Jaeckle KA, Maurer MJ, Reid JM, Ames MM, Hardwick JS, et al. Phase ii trial of vorinostat in recurrent glioblastoma multiforme: A north central cancer treatment group study. Journal of clinical oncology. 2009;27:2052.

30. Yoo CB, Jones PA. Epigenetic therapy of cancer: Past, present and future. Nature reviews Drug discovery. 2006;5:37–50.

31. Wolfson W. Epigenetic cancer therapies emerge out of the lab into the limelight. Chemistry & biology. 2013;20:455–6.

32. Wang M, Lin H. Understanding the function of mammalian sirtuins and protein lysine acylation. Annual Review of Biochemistry. 2021;90.

33. Chai X, Guo J, Dong R, Yang X, Deng C, Wei C, et al. Quantitative acetylome analysis reveals histone modifications that may predict prognosis in hepatitis b-related hepatocellular carcinoma. Clinical and translational medicine. 2021;11:e313.

34. Bolden JE, Peart MJ, Johnstone RW. Anticancer activities of histone deacetylase inhibitors. Nature reviews Drug discovery. 2006;5:769–84.

35. Kelly WK, Marks PA. Drug insight: Histone deacetylase inhibitors—development of the new targeted anticancer agent suberoylanilide hydroxamic acid. Nature Clinical Practice Oncology. 2005;2:150–7.

36. Richon VM, Sandhoff TW, Rifkind RA, Marks PA. Histone deacetylase inhibitor selectively induces p21WAF1 expression and gene-associated histone acetylation. Proceedings of the National Academy of Sciences. 2000;97:10014–9.

37. Yoo C, Ryu M-H, Na Y-S, Ryoo B-Y, Lee C-W, Kang Y-K. Vorinostat in combination with capecitabine plus cisplatin as a first-line chemotherapy for patients with metastatic or unresectable gastric cancer: Phase ii study and biomarker analysis. British journal of cancer. 2016;114:1185–90.

38. Rana Z, Diermeier S, Hanif M, Rosengren RJ. Understanding failure and improving treatment using hdac inhibitors for prostate cancer. Biomedicines. 2020;8:22.

39. Falkenberg KJ, Gould CM, Johnstone RW, Simpson KJ. Genome-wide functional genomic and transcriptomic analyses for genes regulating sensitivity to vorinostat. 2014;1.

40. Falkenberg KJ, Newbold A, Gould CM, Luu J, Trapani JA, Matthews GM, et al. A genome scale {RNAi} screen identifies {GLI1} as a novel gene regulating vorinostat sensitivity. 2016;23:1209–18.

41. O’Neil NJ, Bailey ML, Hieter P. Synthetic lethality and cancer. Nat Rev Genet. 2017;18:613–23.

42. Marigo V, Johnson RL, Vortkamp A, Tabin CJ. Sonic hedgehog differentially regulates expression of gli and gli3 during limb development. Developmental biology. 1996;180:273–83.

43. Skoda AM, Simovic D, Karin V, Kardum V, Vranic S, Serman L. The role of the hedgehog signaling pathway in cancer: A comprehensive review. Bosnian journal of basic medical sciences. 2018;18:8.

44. Niyaz M, Khan MS, Mudassar S. Hedgehog signaling: An achilles’ heel in cancer. Translational oncology. 2019;12:1334–44.

45. Edgar R, Domrachev M, Lash AE. Gene expression omnibus: NCBI gene expression and hybridization array data repository. Nucleic acids research. 2002;30:207–10.

46. Barrett T, Wilhite SE, Ledoux P, Evangelista C, Kim IF, Tomashevsky M, et al. NCBI geo: Archive for functional genomics data sets—update. Nucleic acids research. 2012;41:D991–5.

47. Leinonen R, Sugawara H, Shumway M, on behalf of the International Nucleotide Sequence Database Collaboration. The Sequence Read Archive. 2011;39:D19–D21.

48. Kodama Y, Shumway M, Leinonen R, on behalf of the International Nucleotide Sequence Database Collaboration. The sequence read archive: explosive growth of sequencing data. 2012;40:D54–D56.

49. Hamosh A, Scott AF, Amberger JS, Bocchini CA, McKusick VA. Online mendelian inheritance in man (omim), a knowledgebase of human genes and genetic disorders. Nucleic acids research. 2005;33 suppl_1:D514–7.

50. Amberger JS, Bocchini CA, Scott AF, Hamosh A. OMIM. org: Leveraging knowledge across phenotype–gene relationships. Nucleic acids research. 2019;47:D1038–43.

51. Isella C, Cantini L, Bellomo SE, Medico E. TCGAcrcmRNA: TCGA crc 450 mRNA dataset. 2020.

52. Reimers M, Carey VJ. [8] Bioconductor: An Open Source Framework for Bioinformatics and Computational Biology. 2006;119–34.

53. Network CGA, others. Comprehensive molecular characterization of human colon and rectal cancer. Nature. 2012;487:330.

54. Grossman RL, Heath AP, Ferretti V, Varmus HE, Lowy DR, Kibbe WA, et al. Toward a shared vision for cancer genomic data. New England Journal of Medicine. 2016;375:1109–12.

55. Printz C. Genomic data commons ushers in new era for information sharing. Cancer. 2016;122:2777–8.

56. Jensen MA, Ferretti V, Grossman RL, Staudt LM. The nci genomic data commons as an engine for precision medicine. Blood. 2017;130:453–9.

57. Zhang Z, Hernandez K, Savage J, Li S, Miller D, Agrawal S, et al. Uniform genomic data analysis in the nci genomic data commons. Nature communications. 2021;12:1–11.

58. Kassambara A. fastqcr: Quality Control of Sequencing Data. {R} package version 0.1.2. 2019.

59. R Core Team. R: A language and environment for statistical computing. Vienna, Austria: R Foundation for Statistical Computing; 2021. https://www.R-project.org/.

60. Souza W de, Souza W de, de Sá Carvalho B, Lopes-Cendes I. Rqc: A Bioconductor Package for Quality Control of {High-Throughput} Sequencing Data. 2018;87.

61. Andrews S. {FastQC}: A quality control tool for high throughput sequence data.

62. Patro R, Duggal G, Love MI, Irizarry RA, Kingsford C. Salmon provides fast and bias-aware quantification of transcript expression. Nat Methods. 2017;14:417–9.

63. Patro R, Mount SM, Kingsford C. Sailfish enables alignment-free isoform quantification from {RNA-seq} reads using lightweight algorithms. Nat Biotechnol. 2014;32:462–4.

64. Bray NL, Pimentel H, Melsted P, Pachter L. Near-optimal probabilistic {RNA-seq} quantification. Nat Biotechnol. 2016;34:525–7.

65. Wu DC, Yao J, Ho KS, Lambowitz AM, Wilke CO. Limitations of alignment-free tools in total {RNA-seq} quantification. BMC Genomics. 2018;19:510.

66. Kim D, Pertea G, Trapnell C, Pimentel H, Kelley R, Salzberg SL. {TopHat2}: accurate alignment of transcriptomes in the presence of insertions, deletions and gene fusions. Genome Biol. 2013;14:R36.

67. Tŕapnell C, Pachter L, Salzberg SL. {TopHat}: discovering splice junctions with {RNA-Seq}. Bioinformatics. 2009;25:1105–11.

68. Trapnell C, Williams BA, Pertea G, Mortazavi A, Kwan G, Baren MJ van, et al. Transcript assembly and quantification by {RNA-Seq} reveals unannotated transcripts and isoform switching during cell differentiation. Nat Biotechnol. 2010;28:511–5.

69. Langmead B, Trapnell C, Pop M, Salzberg SL. Ultrafast and memory-efficient alignment of short {DNA} sequences to the human genome. Genome Biol. 2009;10:R25.

70. iGenomes {Ready-To-Use} Reference Sequences and Annotations.

71. The SAM/BAM Format Specification Working Group. Sequence {Alignment/Map} Format Specification.

72. Liao Y, Smyth GK, Shi W. The {R} package Rsubread is easier, faster, cheaper and better for alignment and quantification of {RNA} sequencing reads. Nucleic Acids Research. 2019;47:e47.

73. Love MI, Huber W, Anders S. Moderated estimation of fold change and dispersion for {RNA-seq} data with {DESeq2}. Genome Biol. 2014;15:550.

74. Liu H, Motoda H. Feature selection for knowledge discovery and data mining. 1998.

75. Liu H, Motoda H. Computational methods of feature selection. CRC Press; 2007.

76. Guyon I, Elisseeff A. An introduction to variable and feature selection. J Mach Learn Res. 2003;3:1157–82.

77. Kohavi R, John GH. Wrappers for feature subset selection. Artificial Intelligence. 1997;97:273–324.

78. Blum AL, Langley P. Selection of relevant features and examples in machine learning. Artificial Intelligence. 1997;97:245–71.

79. Yu L, Liu H. Efficient feature selection via analysis of relevance and redundancy. J Mach Learn Res. 2004;5:1205–24.

80. Lopes FM, Martins-Jr DC, Barrera J, Cesar-Jr RM. An iterative feature selection method for GRNs inference by exploring topological properties. arXiv:11075000v1. 2011.

81. Anders S, Huber W. Differential expression analysis for sequence count data. Nature Precedings. 2010;1–1.

82. Love MI, Huber W, Anders S. Moderated estimation of fold change and dispersion for rna-seq data with deseq2. Genome biology. 2014;15:1–21.

83. Sun S, Hood M, Scott L, Peng Q, Mukherjee S, Tung J, et al. Differential expression analysis for rnaseq using poisson mixed models. Nucleic acids research. 2017;45:e106–6.

84. Liu H, Zhang F, Mishra SK, Zhou S, Zheng J. Knowledge-guided fuzzy logic modeling to infer cellular signaling networks from proteomic data. Scientific reports. 2016;6:1–12.

85. Mering C von, Huynen M, Jaeggi D, Schmidt S, Bork P, Snel B. STRING: A database of predicted functional associations between proteins. Nucleic acids research. 2003;31:258–61.

86. Szklarczyk D, Franceschini A, Kuhn M, Simonovic M, Roth A, Minguez P, et al. The string database in 2011: Functional interaction networks of proteins, globally integrated and scored. Nucleic acids research. 2010;39 suppl_1:D561–8.

87. Szklarczyk D, Gable AL, Nastou KC, Lyon D, Kirsch R, Pyysalo S, et al. The string database in 2021: Customizable protein–protein networks, and functional characterization of user-uploaded gene/measurement sets. Nucleic Acids Research. 2021;49:D605–12.

88. Fitch JP, Sokhansanj B. Genomic engineering: Moving beyond DNA sequence to function. Proceedings of the IEEE. 2000;88:1949–71.

89. Liang S, Fuhrman S, Somogyi R. Reveal, a general reverse engineering algorithm for inference of genetic network architectures. Pac Symp Biocomput. 1998;18–29.

90. Glass L, Kauffman SA. The logical analysis of continuous, non-linear biochemical control networks. J Theor Biol. 1973;39:103–29.

91. Tegner J, Yeung MKS, Hasty J, Collins JJ. Reverse engineering gene networks: Integrating genetic perturbations with dynamical modeling. Proc Natl Acad Sci U S A. 2003;100:5944–9.

92. Sokhansanj BA, Fitch JP, Quong JN, Quong AA. Linear fuzzy gene network models obtained from microarray data by exhaustive search. BMC Bioinformatics. 2004;5:108.

93. Woolf PJ, Wang Y. A fuzzy logic approach to analyzing gene expression data. Physiol Genomics. 2000;3:9–15.

94. Raza K. Fuzzy logic based approaches for gene regulatory network inference. Artif Intell Med. 2019;97:189–203.

95. Gormley M, Akella VU, Quong JN, Quong AA. An integrated framework to model cellular phenotype as a component of biochemical networks. Adv Bioinformatics. 2011;2011:608295.

96. Raza K. Fuzzy logic based approaches for gene regulatory network inference. Artif Intell Med. 2019;97:189–203.

97. Sokhansanj BA, Garnham JB, Patrick Fitch J. Interpreting microarray data to build models of microbial genetic regulation networks. 2002. doi:10.1117/12.469450.

98. Sokhansanj BA, Fitch JP. {URC} fuzzy modeling and simulation of gene regulation.

99. Gormley M, Akella VU, Quong JN, Quong AA. An integrated framework to model cellular phenotype as a component of biochemical networks. Adv Bioinformatics. 2011;2011:608295.

100. Datta S, Sokhansanj BA. Accelerated search for biomolecular network models to interpret high-throughput experimental data. BMC Bioinformatics. 2007;8:258.

101. Combs WE, Andrews JE. Combinatorial rule explosion eliminated by a fuzzy rule configuration. 1998;6:1–11.

102. Zadeh LA. Fuzzy sets. Information and Control. 1965;8:338–53. doi:10.1016/s0019-9958(65)90241-x.

103. Mendel JM. Fuzzy logic systems for engineering: a tutorial. 1995;83:345–77.

104. Sokhansanj BA, Fitch JP, Quong JN, Quong AA. Linear fuzzy gene network models obtained from microarray data by exhaustive search. BMC Bioinformatics. 2004;5:108.

105. Anderson C, Ray W. Improved maximum likelihood estimators for the gamma distribution. Communications in Statistics-Theory and Methods. 1975;4:437–48.

106. Forbes C, Evans M, Hastings N, Peacock B. Statistical distributions. John Wiley & Sons; 2011.

107. Kulkarni H, Powar S. A new method for interval estimation of the mean of the gamma distribution. Lifetime data analysis. 2010;16:431–47.

108. Singh A, Singh AK, Iaci RJ. Estimation of the exposure point concentration term using a gamma distribution. In: In usepa, ed. Citeseer; 2002.

109. Ahrens JH, Dieter U. Generating gamma variates by a modified rejection technique. Communications of the ACM. 1982;25:47–54.

110. Ahrens JH, Dieter U. Computer methods for sampling from gamma, beta, poisson and bionomial distributions. Computing. 1974;12:223–46.

111. Zhang J, Zhu W, Wang Q, Gu J, Huang LF, Sun X. Differential regulatory network-based quantification and prioritization of key genes underlying cancer drug resistance based on time-course rna-seq data. PLoS computational biology. 2019;15:e1007435.

112. Teschendorff AE, Severini S. Increased entropy of signal transduction in the cancer metastasis phenotype. BMC systems biology. 2010;4:1–15.

113. Ortiz-Arroyo D, Hussain DA. An information theory approach to identify sets of key players. In: European conference on intelligence and security informatics. Springer; 2008. pp. 15–26.

114. Jalili M, Salehzadeh-Yazdi A, Asgari Y, Arab SS, Yaghmaie M, Ghavamzadeh A, et al. CentiServer: A comprehensive resource, web-based application and r package for centrality analysis. PloS one. 2015;10:e0143111.

115. Shannon C. A mathematical theory of communication, bell systems technol. J. 1948;27:379–423.

116. Kaplan EL, Meier P. Nonparametric estimation from incomplete observations. Journal of the American statistical association. 1958;53:457–81.

117. Goel MK, Khanna P, Kishore J. Understanding survival analysis: Kaplan-meier estimate. International journal of Ayurveda research. 2010;1:274.

118. Rich JT, Neely JG, Paniello RC, Voelker CC, Nussenbaum B, Wang EW. A practical guide to understanding kaplan-meier curves. Otolaryngology—Head and Neck Surgery. 2010;143:331–6.

119. Cox DR. Regression models and life-tables. Journal of the Royal Statistical Society: Series B (Methodological). 1972;34:187–202.

120. Bland JM, Altman DG. The logrank test. Bmj. 2004;328:1073.

121. Kruzelock RP, Short W. Colorectal cancer therapeutics and the challenges of applied pharmacogenomics. Current problems in cancer. 2007;31:315–66.

122. Lemery S, Keegan P, Pazdur R. First fda approval agnostic of cancer site-when a biomarker defines the indication. The New England journal of medicine. 2017;377:1409–12.

123. Board PATE. Colon cancer treatment (pdq): Health professional version. PDQ Cancer Information Summaries, originally published by the National Cancer Institute [Internet]. 2021.

124. Llosa NJ, Cruise M, Tam A, Wicks EC, Hechenbleikner EM, Taube JM, et al. The vigorous immune microenvironment of microsatellite instable colon cancer is balanced by multiple counter-inhibitory checkpoints. Cancer discovery. 2015;5:43–51.

125. Dudley JC, Lin M-T, Le DT, Eshleman JR. Microsatellite instability as a biomarker for pd-1 blockade. Clinical Cancer Research. 2016;22:813–20.

126. LaBaer J, Garrett MD, Stevenson LF, Slingerland JM, Sandhu C, Chou HS, et al. New functional activities for the p21 family of cdk inhibitors. Genes & development. 1997;11:847–62.

127. Johnstone RW, Licht JD. Histone deacetylase inhibitors in cancer therapy: Is transcription the primary target? Cancer cell. 2003;4:13–8.

128. Marks PA, Richon VM, Miller T, Kelly WK. Histone deacetylase inhibitors. Advances in cancer research. 2004;91:137–68.

129. Maloverjan A, Piirsoo M, Michelson P, Kogerman P, Østerlund T. Identification of a novel serine/threonine kinase ulk3 as a positive regulator of hedgehog pathway. Experimental cell research. 2010;316:627–37.

130. Murone M, Luoh S-M, Stone D, Li W, Gurney A, Armanini M, et al. Gli regulation by the opposing activities of fused and suppressor of fused. Nature cell biology. 2000;2:310–2.

131. Koyabu Y, Nakata K, Mizugishi K, Aruga J, Mikoshiba K. Physical and functional interactions between zic and gli proteins. Journal of Biological Chemistry. 2001;276:6889–92.

132. Palencia-Campos A, Ullah A, Nevado J, Yildirim R, Unal E, Ciorraga M, et al. GLI1 inactivation is associated with developmental phenotypes overlapping with ellis–van creveld syndrome. Human molecular genetics. 2017;26:4556–71.

133. Lo H-W, Zhu H, Cao X, Aldrich A, Ali-Osman F. A novel splice variant of gli1 that promotes glioblastoma cell migration and invasion. Cancer research. 2009;69:6790–8.

134. Loo LSW, Soetedjo AAP, Lau HH, Ng NHJ, Ghosh S, Nguyen L, et al. BCL-xL/bcl2l1 is a critical anti-apoptotic protein that promotes the survival of differentiating pancreatic cells from human pluripotent stem cells. Cell Death & Disease. 2020;11:1–18.

135. Adams JM, Cory S. The bcl-2 protein family: Arbiters of cell survival. Science. 1998;281:1322–6.

136. Carpenter RL, Lo H-W. Hedgehog pathway and gli1 isoforms in human cancer. Discovery medicine. 2012;13:105.

137. Merchant M, Vajdos FF, Ultsch M, Maun HR, Wendt U, Cannon J, et al. Suppressor of fused regulates gli activity through a dual binding mechanism. Molecular and cellular biology. 2004;24:8627–41.

138. Zhang Y, Fu L, Qi X, Zhang Z, Xia Y, Jia J, et al. Structural insight into the mutual recognition and regulation between suppressor of fused and gli/ci. Nature communications. 2013;4:1–12.

139. Kogerman P, Grimm T, Kogerman L, Krause D, Undén AB, Sandstedt B, et al. Mammalian suppressor-of-fused modulates nuclear–cytoplasmic shuttling of gli-1. Nature cell biology. 1999;1:312–9.

140. Stone DM, Murone M, Luoh S, Ye W, Armanini MP, Gurney A, et al. Characterization of the human suppressor of fused, a negative regulator of the zinc-finger transcription factor gli. Journal of cell science. 1999;112:4437–48.

141. Han H, Cho J-W, Lee S, Yun A, Kim H, Bae D, et al. TRRUST v2: An expanded reference database of human and mouse transcriptional regulatory interactions. Nucleic acids research. 2018;46:D380–6.

142. Agarwal NK, Qu C, Kunkulla K, Liu Y, Vega F. Transcriptional regulation of serine/threonine protein kinase (akt) genes by glioma-associated oncogene homolog 1. Journal of Biological Chemistry. 2013;288:15390–401.

143. Nye MD, Almada LL, Fernandez-Barrena MG, Marks DL, Elsawa SF, Vrabel A, et al. The transcription factor gli1 interacts with smad proteins to modulate transforming growth factor *β*-induced gene expression in a p300/creb-binding protein-associated factor (pcaf)-dependent manner. Journal of Biological Chemistry. 2014;289:15495–506.

144. Ruan H, Luo H, Wang J, Ji X, Zhang Z, Wu J, et al. Smoothened-independent activation of hedgehog signaling by rearranged during transfection promotes neuroblastoma cell proliferation and tumor growth. Biochimica et Biophysica Acta (BBA)-General Subjects. 2016;1860:1961–72.

145. Shahi MH, Lorente A, Castresana JS. Hedgehog signalling in medulloblastoma, glioblastoma and neuroblastoma. Oncology reports. 2008;19:681–8.

146. Gershon TR, Shirazi A, Qin L-X, Gerald WL, Kenney AM, Cheung N-K. Enteric neural crest differentiation in ganglioneuromas implicates hedgehog signaling in peripheral neuroblastic tumor pathogenesis. PloS one. 2009;4:e7491.

147. Zhu T, Zheng J, Zhuo W, Pan P, Li M, Zhang W, et al. ETV4 promotes breast cancer cell stemness by activating glycolysis and cxcr4-mediated sonic hedgehog signaling. Cell Death Discovery. 2021;7:1–15.

148. Mazzone A, Gibbons SJ, Eisenman ST, Strege PR, Zheng T, D’Amato M, et al. Direct repression of anoctamin 1 (ano1) gene transcription by gli proteins. The FASEB Journal. 2019;33:6632–42.

149. Mroz MS, Keely SJ. Epidermal growth factor chronically upregulates ca2+-dependent cl-conductance and tmem16a expression in intestinal epithelial cells. The Journal of physiology. 2012;590:1907–20.

